# RFW captures species-level full profile of metagenomic functions via integrating genome annotation information

**DOI:** 10.1101/2024.03.19.585660

**Authors:** Kai Mi, Xingyin Liu

## Abstract

Functional profiling on whole-metagenome shotgun sequencing (WMS) has made great contribution to the development of our understanding in microbe-host interactions. In this work, we revealed that severe microbial functional information loss of current functional profiling methods existed at both taxon-level and community-level. To correct the distortion brought by information incompleteness, we developed a new framework, RFW (Reference based functional profile inference on WMS), to infer microbial functional abundance on WMS through utilizing information from genome function annotation and WMS taxonomic profile. Furthermore, we built up a new algorithm for absolute abundance change quantification of microbial function between groups under RFW framework. By applying RFW to several datasets related to autism spectrum disorder and colorectal cancer, we revealed that RFW greatly renewed our knowledge in downstream analysis, including differential microbial function identification, association analysis between microbial function and host phenotype, etc. RFW are open-source and freely available at https://github.com/Xingyinliu-Lab/RFW.

## Introduction

Understanding the integral roles of microbial function in maintenance of metabolism and microbiome homeostasis is key to deciphering the host-microbiome interaction[1, 2]. Increasing functional profiling tools for shotgun metagenomic data by quantification of the microbial genes and metabolic pathways have been reported in recent years [3–8].

Whole-metagenome shotgun sequencing (WMS) has empowered species-level resolution of microbial functions. The tools for analyzing WMS functional profile can be classified into two main categories, i.e., assembly-based and sequence-based. An early practice of assembly-based functional profiling was that Qin et al. annotated functions of predicted open reading frames (ORFs) from assembled contigs and featured the human gut microbial gene catalogue[9]. In the same period, sequence-based functional profiling tool MG RAST server[4] was proposed, assigning the potential function to each sequence by BLAST[10] search against the SEED[11] database. Subsequently, MEGAN[5] introduced KEGG[12] metabolic maps for microbial bioactivity pathway analysis. Given the latter sequence-based strategy would be more friendly to functional abundance quantification, most follow-up methods, including ShotMAP[13], COGNIZER[14], FMAP[15], *mi-faser*[16], Fun4Me[17] and HUMAnN[6–8] etc., were constructed upon the strategy. These methods built their pipeline utilizing different alignment programs (BLAST, DIAMOND[18], FragGeneScan[19], or RAPSearch2[20], etc.), different reference database(KEGG, Pfam[21], GO[22], SEED, eggNOG[23] or manually curated database, etc.). Instead of sequence alignment, Carnelian et al. trained a Vowpal Wabbit one-against-all (OAA) classifier to predict the function of k-merized sequence directly[24]; Hoarfrost et al. presented LookingGlass for interpretation of short DNA reads by deep learning[25].

Among these methods, HUMAnN is the state-of-the-art one and has been applied in multiple recent studies[26–30]. Compared with other methods, HUMAnN counted the reads alignment hits in a weighted manner rather than best-hit. Meanwhile, HUMAnN embraced a tiered-search manner in sequence alignment which significantly accelerated the mapping computation and accuracy. HUMAnN were also used as benchmark for genus-level microbial function prediction from 16S rRNA sequencing, e.g., PICRUSt, Tax4Fun and PICRUSt2[31, 32].

Currently, for all shotgun metagenomic functional profiling methods, the information was extracted from the sequencing reads. However, Zhernakova et al. reported that the profiled gene richness had a significant association with sequencing depth, while the microbial composition was robust confronted with sequencing depth[33]. Furthermore, PICRUSt were in better agreement with WMS HUMAnN with deeper sequencing depth[34]. These two reports raised our concern whether WMS was adequate for functional profiling as a proxy for the true metagenome.

Here we present RFW (Reference based functional profile inference on WMS), a framework for functional profile prediction from WMS taxonomic profile using the genome-function reference database. Our results show that HUMAnN in WMS exhibited severe functional information incompleteness due to sampling depth and RFW fixed the issue effectively. Meanwhile, to correct potential bias produced from relative abundance, we developed DFSCA-BC (Differential FSCA(Functional Sub-Community Abundance) Identification with Bias Correction), a new method for differential absolute microbial functional abundance analysis between groups, inherited from our previously published taxonomic absolute abundance change quantification method QMD[35]. Finally, we applied RFW and DFSCA-BC to several datasets to demonstrate the full profile of metagenomic functions.

RFW and DFSCA-BC are open-source and freely available at https://github.com/Xingyinliu-Lab/RFW. RFW supports multiple forms of microbial function description(termed in KEGG Orthology (KO), EC (Enzyme Commission) numbers[36] and Clusters of Orthologous Genes (COG)[37]). Given the EC numbers could be directly mapped into MetaCyc[38, 39] pathways, we selected 4th Level EC enzyme families (abbreviated as enzyme hereafter) as a basis in following analysis and comparisons.

## Results

### Function profiling on WMS exhibited functional information loss

To check whether WMS is sufficient for microbial function profiling, a reference database was constructed by annotating protein functions of high-quality genomes from 65,703 bacterial and archaeal species, representing 317,542 genomes (see Methods), because the microbe genome was considered as the most comprehensive source of information on its bioactivity function.

With the reference database, we first investigated whether HUMAnN was able to identify all microbial functions in WMS on cultured bacterial isolates, in which the potential effects of complex microbiome composition on functional profiling were reduced. The WMS dataset came from Elie et al[40, 41], involving 15 bacteria isolated from C57BL/6J mice (see Supplementary Table 1). With comparing the genome-annotated and HUMAnN reported bacteria enzymes, a discernible lack in WMS functional profiling was observed and *approximately 30% genome functions were mis-detected by* HUMAnN (Figure 1A) in a species-dependent manner (Figure 1B-E). For example, WMS on the *Escherichia coli* isolate covered 88.3% enzymes (962/(962+128)) (Figure 1B), comparatively 50.3% on *Parabacteroides goldsteinii* (366/(366+362)) (Figure 1E). Totally. 109 EC enzymes were missing in more than half of the 15 bacteria, listed in Supplementary Table 2. Of these enzymes, EC 3.1.3.100 (thiamine phosphate phosphatase) and EC 3.6.5.4 (signal-recognition-particle GTPase) were encoded by all 15 bacteria inferred from the corresponding genomes. The enzyme protein information collected from KEGG and UniProt as evidence also proved the bacterial encoding capability (see Supplementary Table 3). However, none WMS functional profiling detected these two enzymes.

**Figure 1.**
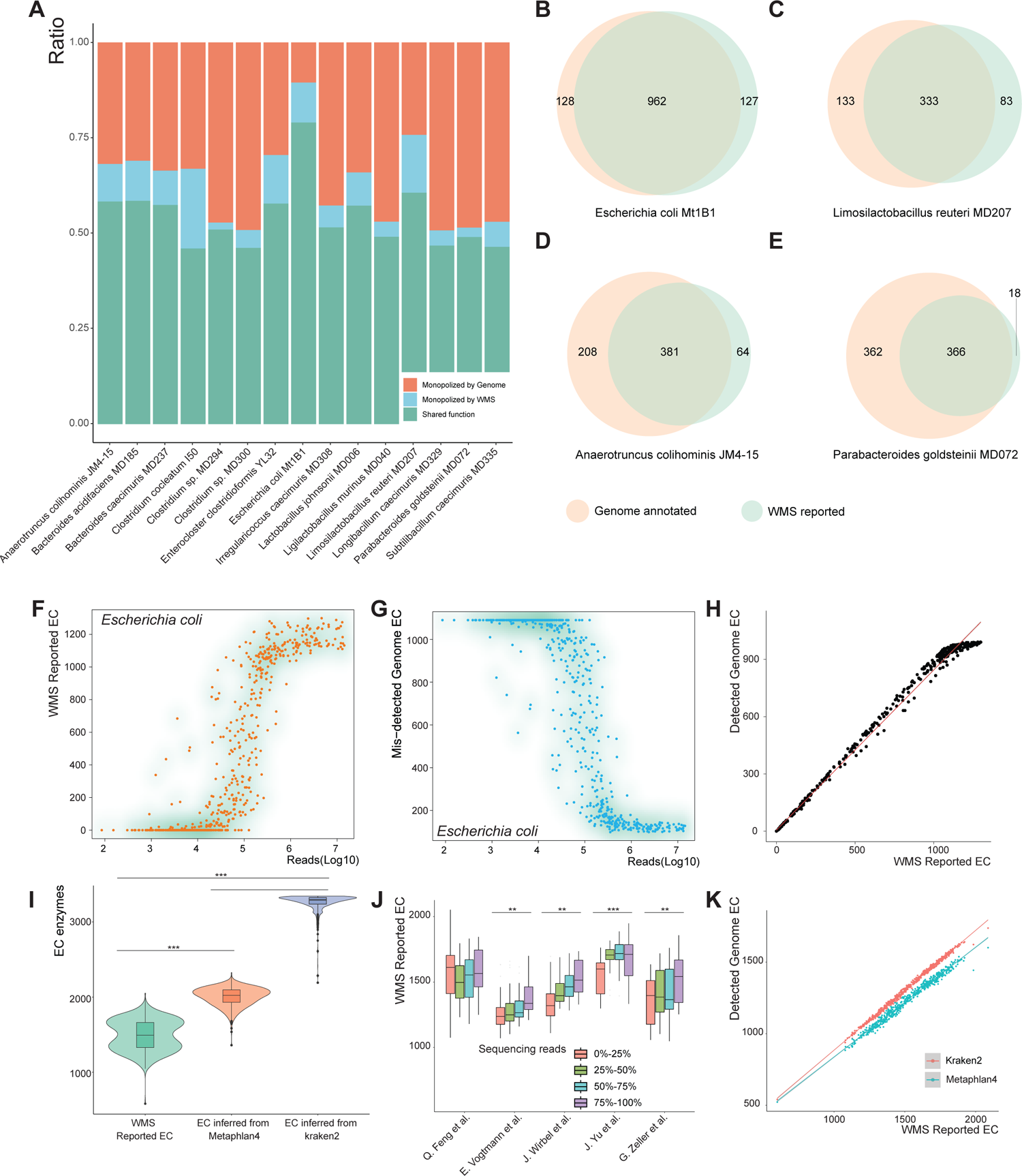
Severe functional information loss in WMS. **A,** Approximately 30% genome functions were lost in HUMAnN analysis of WMS on bacterial isolates. **B-E,** Venn plots of bacterial isolate WMS-HUMAnN reported and genome-annotated enzymes for *Escherichia coli Mt1B1* (**B**), *Limosilactobacillus reuteri MD207* (**C**), *Anaerotruncus colihominis JM4-15* (**D**), and *Parabacteroides goldsteinii MD072* (**E**). **F,** HUMAnN reported enzymes of *Escherichia coli* from WMS on gut microbiome degraded sharply with shallow sequencing depth (by reads). **G,** Shallow sequencing depth missed genome-annotated enzymes of *Escherichia coli* from WMS on gut microbiome. **H,** HUMAnN reported more enzymes means it could cover more genome annotated functions for *Escherichia coli*. **I,** Community-level enzymes reported by WMS-HUMAnN were significantly less than that inferred from microbial taxonomic profile. **J,** The number of community-level enzymes reported by WMS increased with sequencing depth. **K,** Community-level enzymes reported by HUMAnN had a positive linear correlation with the covered genome annotated enzymes. *Student t test, *: P < 0.05, **: P < 0.01, ***: P < 0.001. Detected genome EC: enzymes which were inferred from genome-annotation and also reported by HUMAnN. Mis-Detected genome EC: enzymes which were inferred from genome-annotation and failed to be detected by HUMAnN*.

Next, we enrolled a total of 589 WMS samples from healthy people, Adenoma or Colorectal Cancer patients in five established cohorts, including Feng et al.[42] (ERP008729), Vogtmann et al.[43] (PRJEB12449), Wirbel et al.[44] (PRJEB27928), Yu et al.[45] (PRJEB10878), and Zeller et al.[46] (PRJEB6070) to check HUMAnN performance under complex microbiome composition. Ten common bacteria, including *Escherichia coli*, *Akkermansia muciniphila*, and *Alistipes indistinctus* etc., were picked to evaluate the taxon-level functional profiling of HUMAnN. Relative abundances (calculated by Kraken2[47]) multiplied by WMS reads were used to represent the sequencing depth for each taxon. When sequencing reads aligned to *Escherichia coli* were less than 10^6^, the number of EC enzymes reported by HUMAnN declined sharply with decreasing sequencing reads (Figure 1F). Synchronously, the mis-detected genome function increased with shallow sequencing depth (Figure 1G). It was also noticed that HUMAnN-reported enzyme had a highly inherent consistency with the genome annotated function (Figure 1H). Given the total sequencing reads in a WMS sample was about 1e10^7^ to 2e10^7^ on average (Supplementary Figure 1), hence, WMS-reported functions would be incomplete if the relative abundance of *Escherichia coli* was lower than 5% to 10%. Similar conclusion could be drawn for the remaining nine bacteria (Supplementary Figure 2). Furthermore, we obtained the same conclusion through another relative abundance calculation method, Metaphlan4[48, 49] (Supplementary Figure 3).

At the community-level, we calculated all potential microbial functions by integrating the genome annotation reference database and taxa information identified by Kraken2 or Metaphlan4. Then, we compared the results with HUMAnN. Also, HUMAnN on WMS exhibited severe functional information loss at the community-level (Figure 1L). Inference from Kraken2 profiling produced more enzymes than that from Metaphlan4, because Kraken2 reported significantly more taxa than Metaphlan4. According to the number of sequencing reads, we divided the enrolled WMS samples into four quarters. Similarly, HUMAnN-reported more enzymes with increased sequencing depth in the community-level (Figure 1J). More reported enzymes mean more genome functions detected (Figure 1K).

### Aromatic compound metabolic pathways were remarkably mis-detected by HUMAnN

The true positive rate (TPR) of HUMAnN detection on the genome functions by Kraken2 (Figure 2A) for around 1100 bacteria enzymes approached to 80-100% in the 589 WMS samples, similarly on Metaphlan4 (Figure 2B). TPR of the remained around 2000 enzymes were less than 80%, and especially, TPR of more than 1000 enzymes were below 20%. For example, EC 1.1.1.108 (carnitine 3-dehydrogenase) were predicted positive in more than 580 samples at the community-level according to the genome information, but only reported in 3 samples by HUMAnN, with a TPR at 0.52% (3/580=0.52%). With comparing HUMAnN TPR with the genome functions of inferred from Kraken2 and from Metaphlan4 taxonomic profiling, respectively, we found the prevalence of inferred genome functions was highly consistent (Figure 2C). For 2486 out of 2746 enzymes, the differences of TPR assessed from these two methods were less than 5%, as cyan dotted in Figure 2C.

**Figure 2.**
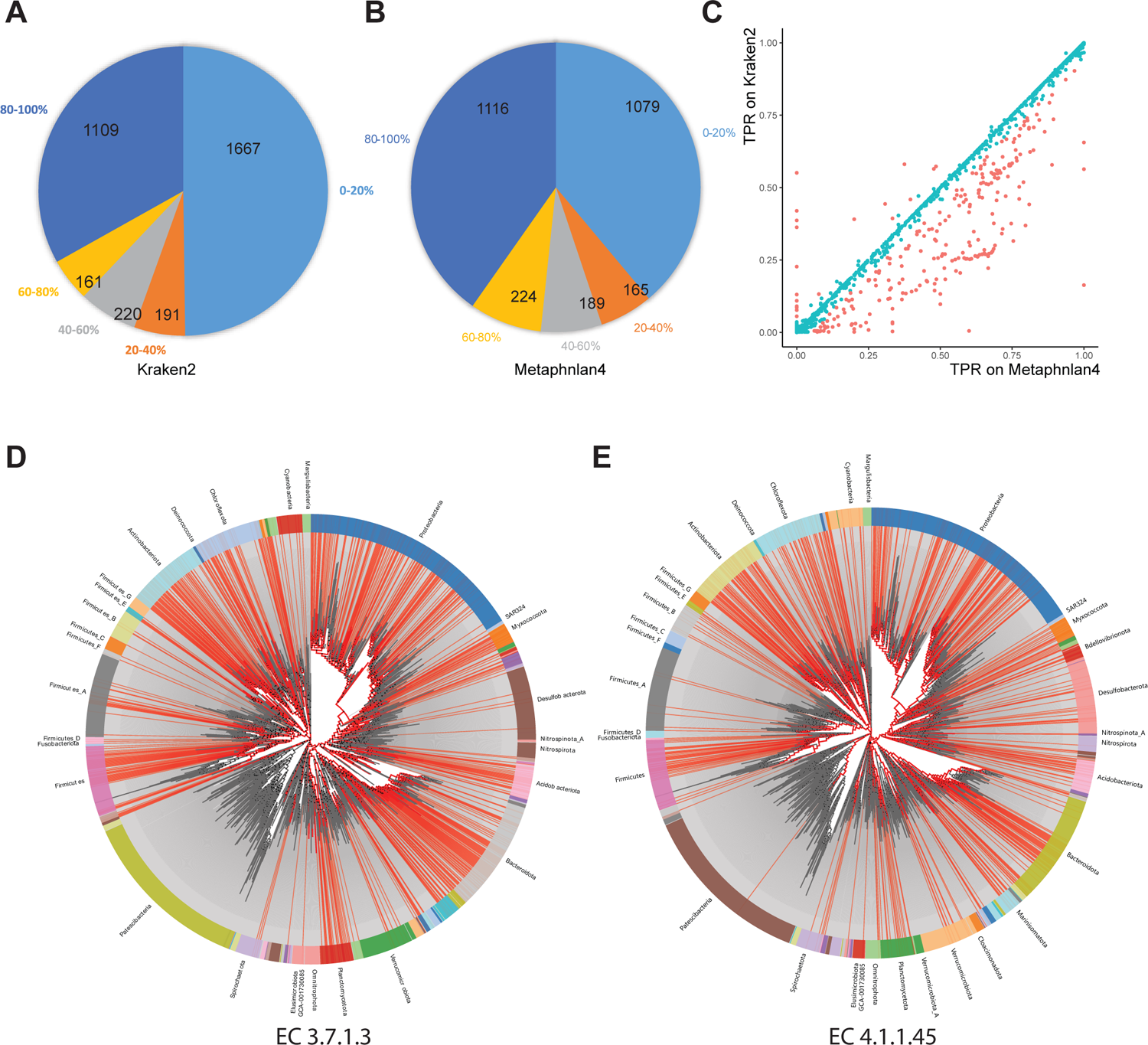
distribution of genome function detection by WMS. **A,** Pie distribution of the true positive rate on Kraken2-assessed genome functions reported by HUMAnN. **B,** Pie distribution of the true positive rate on Metaphlan4-assessed genome functions reported by HUMAnN. **C,** True positive rate on Kraken2-assessed and Metaphlan4-assessed genome functions were highly consistent. **D-E,** Bacteria phylogenetic tree for EC 3.7.1.3 (**D**), and EC 4.1.1.45 (**E**) encoding, red highlight denoted for the bacteria which encode these two enzymes. *0-20%: the number of genome functions with which the proportion of samples that were positively detected by HUMAnN located at 0-20%. 20-40%… etc. hereafter. For example, EC 6.3.2.45 were inferred to be encoded in all 589 samples, but only detected in 444 samples by HUMAnN, then the ratio is 444/589=75.4%. Following this way, we calculated proportions for all genome functions and counted according to the specified interval*.

Next, we made a further survey on WMS function profiling information loss preference. Using fisher exact test, we filtered out enzymes with HUMAnN detection rate significantly changed compared with potential genome function and listed the most involved 20 pathways in Supplementary Table 4 and Supplementary Table 5. These mis-detected enzymes were enriched in the aromatic compound metabolic pathways, including PWY-6954 (superpathway of aromatic compound degradation via 2-hydroxypentadienoate), PWY66-5 (superpathway of cholesterol biosynthesis), ALL-CHORISMATE-PWY (superpathway of chorismate metabolism), and PWY-5655 (L-tryptophan degradation IX) etc.

Take PWY-5655 (L-tryptophan degradation IX) as an example, we detailed the enzyme detection at Supplementary Table 6. Of the 13 enzymes involved in this pathway, EC 1.2.1.10 (acetaldehyde dehydrogenase), EC 3.5.1.9 (arylformamidase), EC 4.1.3.39 (4-hydroxy-2-oxovalerate aldolase) obtained a high TRP larger than 90%. For other enzymes, the TRP was relatively lower. Especially, we highlighted the encoding bacteria for EC 3.7.1.3 (L-kynurenine hydrolase) and EC 4.1.1.45 (aminocarboxymuconate-semialdehyde decarboxylase) at the phylogenetic tree by AnnoTree[50]. AnnoTree stated bacteria from *Actinobacteriota*(24.96%), *Bacteroidota*(18.39%), *Firmicutes*(7.00%), *Proteobacteria*(38.00%) were capable to encode EC 3.7.1.3 (Figure 2D) and bacteria from *Actinobacteriota*(27.93%), *Bacteroidota*(16.32%), *Firmicutes*(7.53%), *Proteobacteria*(31.96%) were capable to encode EC 4.1.1.45 (Figure 2E). In other words, these two enzymes were relatively common functions in bacteria. Our analysis on all community-level potential genome functions also suggested the two enzymes should be positively detected in most samples. While HUMAnN only reported 8 and 0 positive samples for EC 3.7.1.3 and EC 4.1.1.45, respectively, illustrating how HUMAnN lost important microbial functional information. We also noticed the potential positive samples for EC 3.5.99.5 (2-aminomuconate deaminase) assessed from Metaphlan4 was quite fewer than by HUMAnN, implying that the taxonomic profiling by Metaphlan4 was too conservative.

Together, the above analysis indicated that functional profiling in WMS mediated great information loss. Impressively, the annotated genome function reference database could be a good complement for functional profiling.

### The RFW framework for functional profile inference on WMS

Given the microbial functional information loss in WMS HUMAnN analysis would inevitably lead to deficiencies in fully capturing the underlaying interaction of the host and microbiota, we proposed a new method for functional profile inference on WMS, i.e., RFW (Reference based functional profile inference on WMS) (Figure 3A).

**Figure 3.**
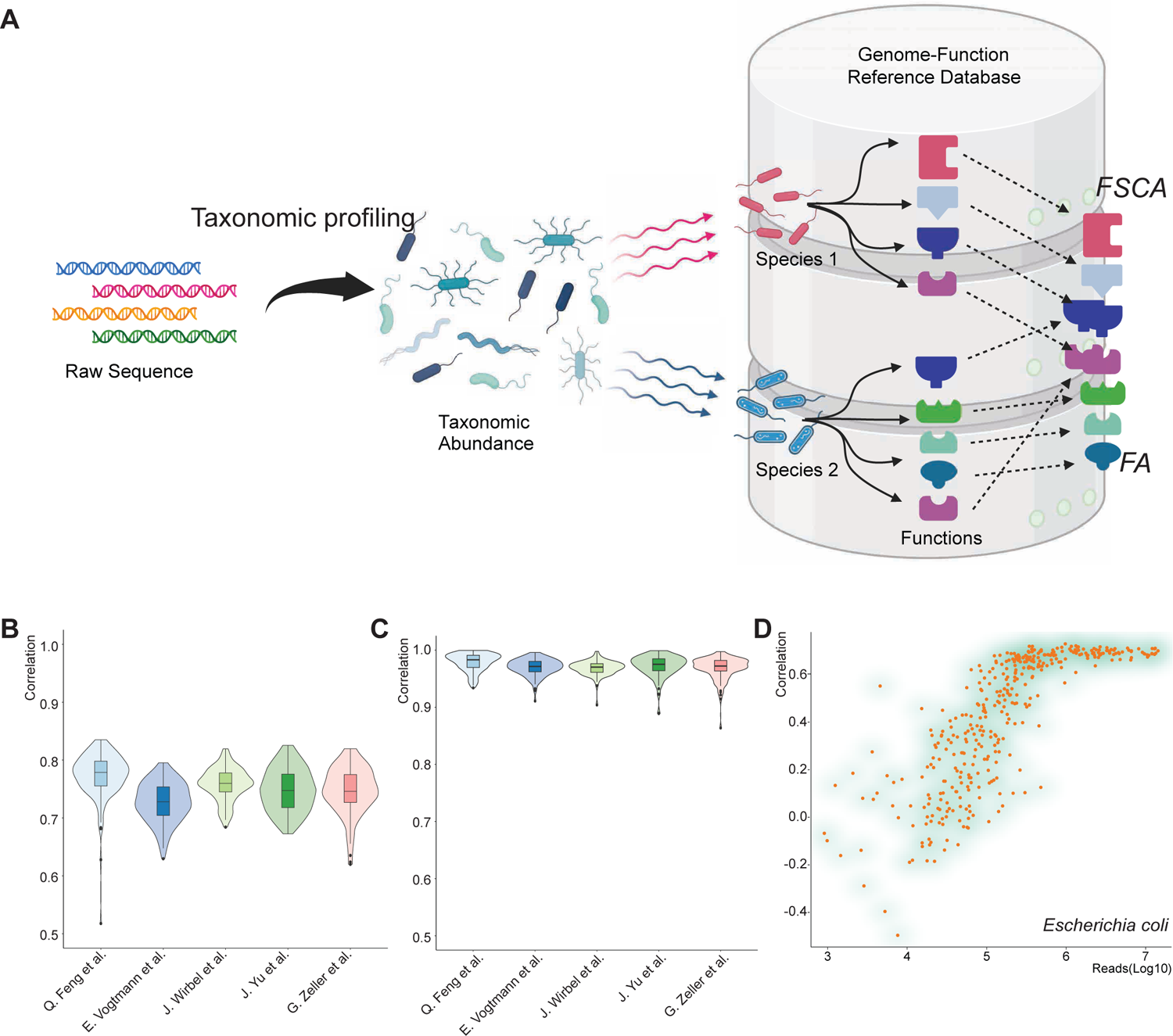
RFW infers microbial function with aid of the genome-function reference database. **A,** Overview on RFW framework. **B,** RFW-FA with taxonomic profiling from Kraken2 had a strong positive Pearson correlation with HUMAnN reported functional abundance. **C,** The correlation between RFW-FA with taxonomic profiling from Kraken2 and that from Metaphlan4 approached to 1. **D,** The sequence depth affected correlations between functional abundance of *Escherichia coli* from RFW with taxonomic profiling from Kraken2 and that from HUMAnN.

To make the inference, denote taxa set identified from taxonomic profiling as *Taxa* = {*Taxon*_1_, *Taxon*_2_, … *Taxon*_*p*_}, where *Taxon*_*i*_ is the ith species in the microbiome. And the corresponding relative abundance of each taxon is (^*x*^*Taxon* ^, *x*^*Taxon* ^, … *x*^*Taxon*)^′^. Denote the genome functions of *Taxon*_*i*_ as {φ^*f*^_Taxon_ |*f* ∈ F}, in which φ^*f*^ = 1 if *Taxon*_*i*_ is capable to encode function *f*, otherwise φ^*f*^ = 0. F represents all potential genome functions of taxa. Denote function counts on *f* of *Taxon*_*i*_ as {ϑ^*f*^ |*f* ∈ F}. The information on genome functions and their counts can be extracted from the reference database.

In RFW framework, we established two schemas for describing microbial functional profile. One is Functional Abundance as FA = {∑_*i*_ *x*_*Taxon*i_ ϑ^*f*^ |*f* ∈ F}, i.e., multiplying the relative abundance of community members by the annotated function counts of their respective reference genomes. Accordingly, *x_Taxon_i* ∗ φ^*f*^*_Taxon_i* ∗ ϑ^*f*^*_Taxon_i* is the taxon-level functional abundance for *Taxon*. Concept of FA was in line with traditional functional profiling methods. In fact, FA was used as the benchmark of mock communities in simulation test of HUMAnN[8, 51] and PICRUSt[34]. We are of the opinion that FA is a middle concept between RNA expression levels and taxon abundance in metagenomics. The second is Functional Sub-Community Abundance as FSCA = {∑_*i*_ *x* |*f* ∈ F}. FSCA was defined as the total relative abundance of sub-community members which could encode the specific function *f*. Hence, FSCA could be easily extended to account the Sub-Community abundance involved into pathways. Besides, FSCA had a powerful advantage in identifying differences in microbial functions between groups.

As expected, RFW-FA with taxonomic profiling from Kraken2 (Figure 3B) had a strong positive Pearson correlation for common functions with HUMAnN, similarly on Metaphlan4 (Supplementary Figure 4). Meanwhile, the correlation between RFW-FA with taxonomic profiling from Kraken2 and that from Metaphlan4 approached to 1, implying that the functional abundance inferred by RFW was robust confronted with different taxonomic profiling methods (Figure 3C). RFW also supports taxon-level functional analysis. Likely, shallow sequence depth reduced correlations between functional abundance of *Escherichia coli* from RFW and that from HUMAnN (Figure 3D). In other words, information loss of HUMAnN in WMS included not only mis-detected microbial functions, but also distorted functional abundance. Analysis on RFW taxon-level abundance for another nine bacteria with taxonomic profiling from Kraken2 (Supplementary Figure 5) supported this deduction, similarly on Metaphlan4 (Supplementary Figure 6).

### The DFSCA-BC algorithm for absolute abundance change quantification of microbial function

Relative abundance is widely used in describing microbiome composition. In contrast, absolute abundance measures the microbial load per unit sample mass (e.g. volume or weight). Our previous work QMD(quantification of microbial absolute abundance differences) has shown a deviation exists between taxon relative abundance changes and absolute abundance changes, moreover, the deviation is exactly equal to the total microbial absolute abundance change between groups[35]. RFW-FSCA measures the relative abundance of taxa encoding some certain microbial function *f* and we found that the deviation still exists in comparing FSCA between groups.

Denote sub-community ℒ_*f*_ for microbial function *f*, i.e., ℒ = {*Taxon*|φ = 1}. The absolute abundance change Δ_*f*_ of sub-community ℒ_*f*_ between Control (*C*) and Treat (*T*) could be depicted as:

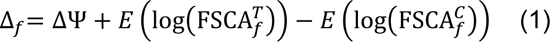

where ΔΨ is the log fold change of total microbial absolute abundance of all taxa between groups, *E* (log(FSCA^*C*^)) and *E* (log(FSCA^*T*^)) are the expectation of logged microbial relative abundance of sub-community ℒ_*f*_, i.e., FSCA_*f*_, of Control and Treat, correspondingly. The deviation ΔΨ can be obtained following the pipeline shown in our previous work QMD[35]. Detailed proof on formula 1 was provided in Supplementary information 1.

Hence, a new statistical hypothesis could be established for identifying absolute abundance changed microbial functions, as

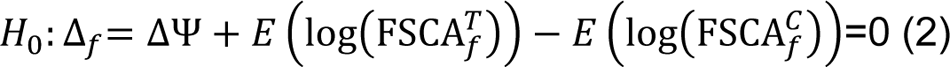

i.e.:

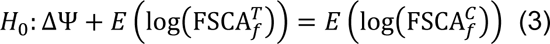

In this way, we built up a new method for differential microbial function identification. We name the hypothesis test as DFSCA-BC (Differential FSCA Identification with Bias Correction). Furthermore, DFSCA-BC can also be used to identify absolute abundance changes of sub-community involved into pathways. Specially, the formula 3 would degrade to *H*_0_: ΔΨ = 0 for microbial functions which are expressed universally in all microbiota, and absolute abundance of these microbial functions would be judged as statistically changed if ΔΨ ≠ 0.

### Completeness of microbial functional profile contributed to higher identification power in differential microbial functional abundance analysis

We applied RFW in identification of gut microbiome function changes between healthy and CRC people as an illustration of benefit from complete functional profiling. To make a fair comparison, DESeq2[52] was performed on RFW-FA and HUMAnN since the definition on Functional Abundance FA in RFW was in line with HUMAnN. The DESeq2 analysis showed significant inconsistencies between RFW and HUMAnN (Figure 4A-E). For example, in the cohort from Vogtmann et al. there were 177 differential microbial functions between Health and CRC under RFW framework, but 32 under HUMAnN, of which only two functions were considered as differential ones by both HUMAnN and RFW (Figure 4B).

**Figure 4.**
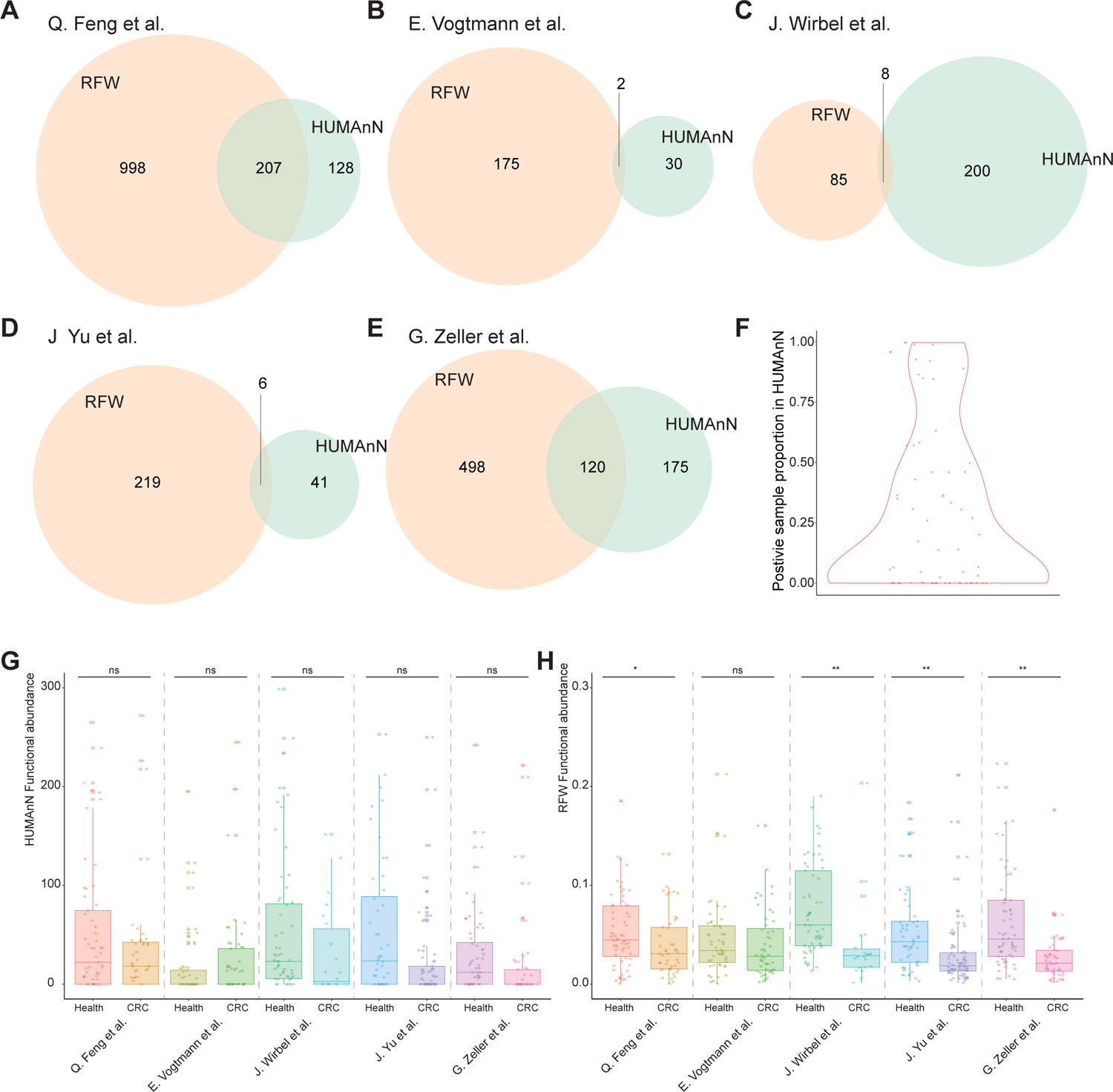
identification of microbial function changes benefited from information completeness of RFW. **A-E,** Venn plot on health and CRC microbial function changes identification results by Deseq2 on RFW-FA and HUMAnN on different cohorts, i.e., Q. Feng et al. (**A**), E. Vogtmann et al. (**B**), J. Wirbel et al. (**C**), J. Yu et al. (**D**), and G. Zeller et al. (**E**), only differential functions with adjusted pvalue <0.05 were counted. **F,** Proportion of positive samples reported by HUMAnN of which the microbial functions were judged as differential ones by Deseq2 on RFW-FA but not on HUMAnN in at least three cohorts. **G-H,** Microbial function EC 2.7.10.1 expression level on different cohorts reported by HUMAnN(**G**) and RFW-FA(**H**). *Deseq2 test, *: adjusted pvalue < 0.05, **: adjusted pvalue < 0.01, ***: adjusted pvalue < 0.001*.

Supplementary Table 7 listed differential microbial functions between Health and CRC identified by RFW-FA but not HUMAnN in at least four cohorts., HUMAnN lost all information on the three enzymes, EC 2.4.1.152, EC 2.4.1.65 and EC 2.1.1.137 among the identified functions. EC 2.4.1.152 and EC 2.4.1.65 participate in GDP-β-L-fucose metabolism and are isoenzyme of FUT family protein in Homo sapiens of which FUT3, FUT6, and FUT9 play important roles in colorectal cancer progression[53], tumor metastasis[54] and cancer cell reprogramming[55]. Gut microbiome derived EC 2.1.1.137 (arsM) and its Homo sapiens isoenzyme AS3MT were both required for full protection against arsenic toxicity[56]. The AS3MT expression were also reported significantly reduced in CRC patients by Gepia[57]. Other microbial functions were partially positive-detected by HUMAnN. For example, EC 2.7.10.1, whose Homo sapiens isoenzyme AATK was frequently hypermethylated [58] and reduced cell proliferation[59] in multi human cancer diseases, was reported in 63% (373/589) samples by HUMAnN.

To illustrate how HUMAnN detection failure affected differential microbial function identification, we summarized the positive proportion of samples in HUMAnN for newly detected differential microbial functions by RFW on at least three cohorts (Figure 4F). It could be seen most proportions was less than 75% by HUMAnN. Again, we took EC 2.7.10.1 as an example, no significant change was found between Health and CRC by HUMAnN (Figure 4G). However, this microbial function was judged to be differential at four cohorts by RFW (Figure 4H). We observed the same comparison tendency on the upper bound of boxplots in Figure 4G and 4H when we examined the functional abundances in the plots. However, the lower bound of boxplots was dragged to nearly 0 in Figure 4G because of information loss of HUMAnN. This is the reason why DESeq2 had difficulty in detecting this enzyme on HUMAnN functional abundance.

Furthermore, we quantified absolute abundance changes of microbial function between Health and CRC by DFSCA-BC algorithm. Significant correlations were observed for fold changes of absolute microbial functions abundance between Health and CRC from different cohorts (Figure 5A). Contrarily, fold changes of taxa abundance from different cohorts showed nearly no correlation (Figure 5B), implying that targeting microbial function might be a more efficient way than targeting microbes in CRC therapy. Figure 5C presented microbial functions with differential abundance between Health and CRC in at least four cohorts by DFSCA-BC. EC 2.4.1.152 and EC 2.4.1.65 were still included in the top differential ones. For the third enzyme, EC 3.2.1.73 catalyzes the hydrolysis linkages in β-D-glucans. β-*D*-glucans were thought to be involved in enhancing immune system against tumor cells and inhibiting tumor invasion and progression[60, 61]. On the other hands, although EC 3.2.1.73 were commonly expressed in various bacteria (Figure 5D), we found that *Bacteroides fragilis* abundance was highly correlated with EC 3.2.1.73 abundance (Figure 5E). Multi researches have supported *Bacteroides fragilis* as a possible etiological candidate for CRC[62, 63]. Together, the information suggested the necessity of investigation on the taxon contribution to microbial function in interpretation of differential microbial functions.

**Figure 5.**
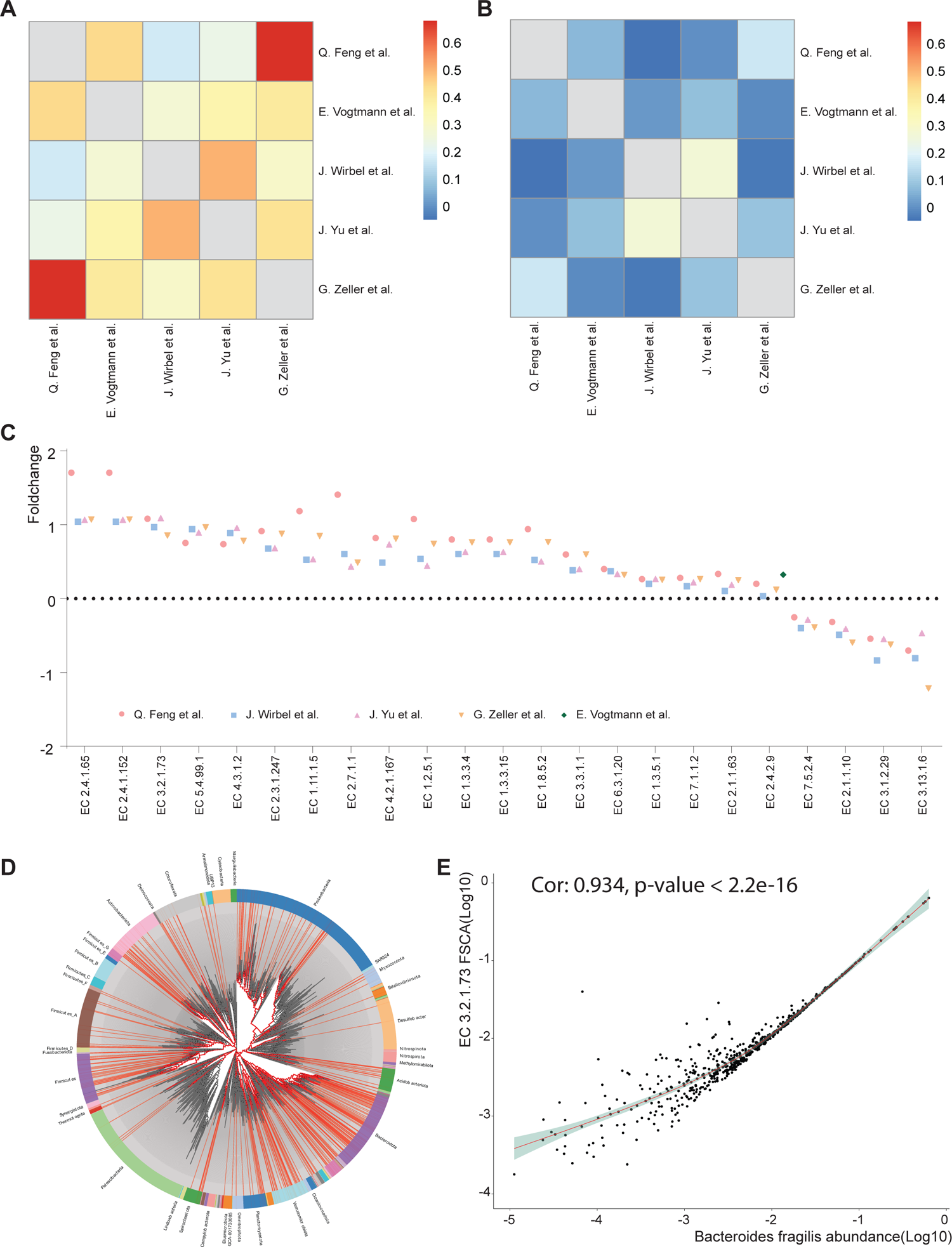
conservation in microbial function changes between Health and CRC. **A,** Correlation of absolute abundance foldchanges of microbial functions between Health and CRC from different cohorts. **B,** Correlation of taxa absolute abundance foldchanges between Health and CRC from different cohorts. **C,** Absolute abundance foldchange of microbial functions which were differential between Health and CRC in at least four cohorts by DFSCA-BC. **D,** Bacteria phylogenetic tree for EC 3.2.1.73 encoding, red highlight denoted for the bacteria which encode these two enzymes. **E,** Bacteroides fragilis abundance was highly correlated with EC 3.2.1.73 abundance.

### RFW enlightened us more insights into associations between microbial functions and host phenotypes

Association analysis is another important scenario in host-microbiome interaction deciphering. To demonstrate the benefit of full-view profile derived from RFW in association analysis, we calculated the Pearson correlation between the microbial functions and ASD symptom severity of 39 ASD children collected from Zhang et al[64]. Evidence has been growing to support associations between autism spectrum disorder (ASD) and gut microbiome dysbiosis[65, 66]. In our analysis, 16 microbial functions were found strongly correlated with different ASD symptom rating criteria (Supplementary Table 8).

With integration of biochemical reaction information from KEGG, Metacyc, Brenda[67] and MetaNetX[68], we found some of the 16 microbial functions were potentially connected by common metabolic pathways, forming metabolic networks. For example, EC 6.3.2.2 and EC 4.2.1.22 were involved in glutamic acid and homocysteine related metabolic network (Figure 6A). Glutamic acid is a precursor of gamma-aminobutyric acid (GABA). Higher blood glutamic acid levels in ASD[69] and its correlation with the severity of social symptoms[70] have been reported. Mutations on Homo sapiens isoenzyme of EC 4.2.1.22, Cystathionine β-synthase (CBS), would lead to homocystinuria and a variety of CNS disturbances, including ASD[71]. Microbial derived EC 4.2.1.22 had a negative correlation with the RBSR (Repetitive Behavior Scale– Revised) score (Figure 6B). HUMAnN failed to detect EC 4.2.1.22 in most samples, thus missed this correlation (Figure 6C). Glutathione cysteine ligase (GCL), the isoenzyme of EC 6.3.2.2 were also found impaired in autism[72]. RFW (Figure 6D) reported a stronger correlation between EC 6.3.2.2 and CARS (Childhood Autism Rating Scale) than HUMAnN (Figure 6E).

**Figure 6.**
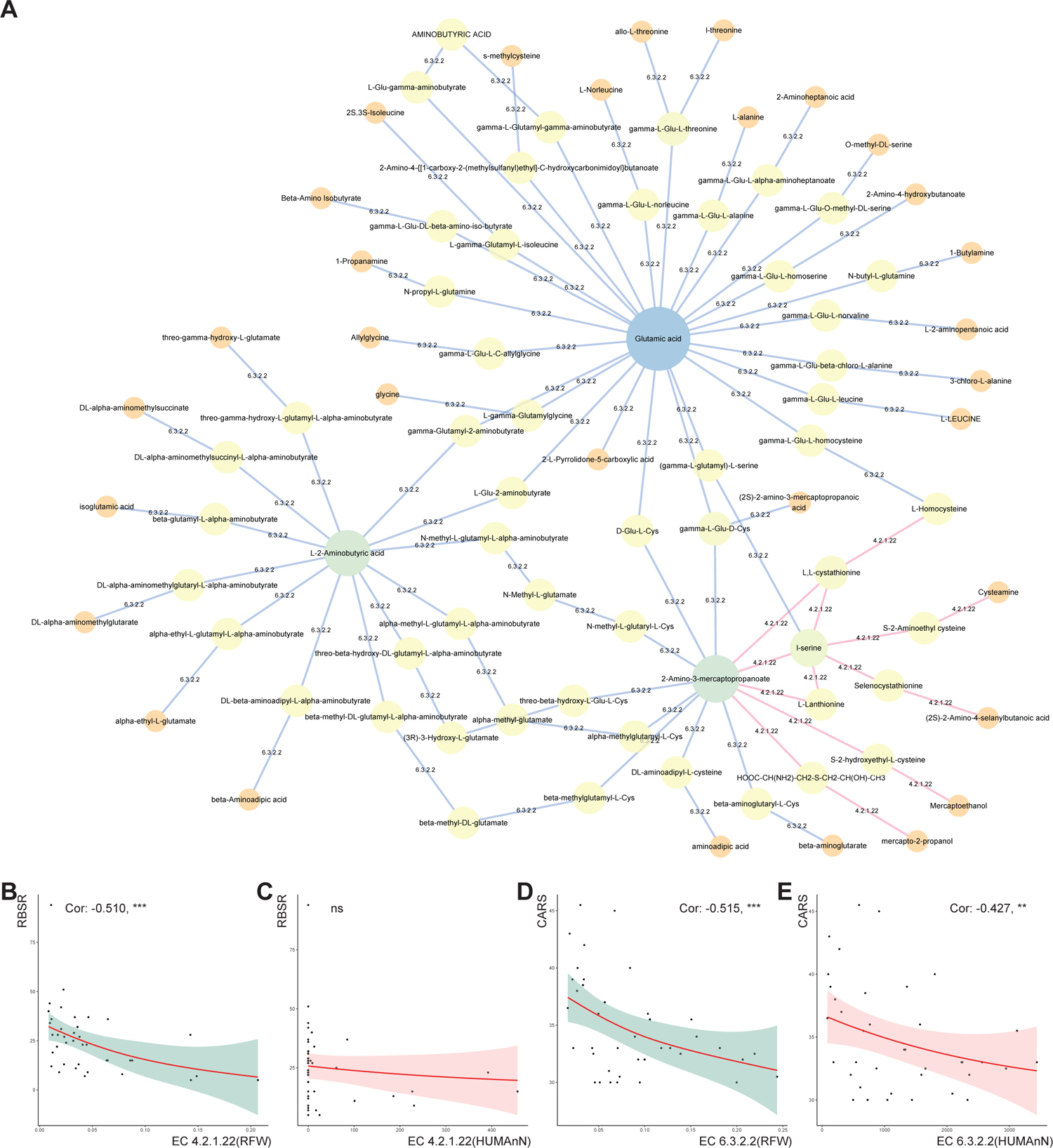
microbial RFW FSCA correlated with ASD symptom. **A,** EC 6.3.2.2 and EC 4.2.1.22 involved metabolic network. **B,** Correlation of EC 4.2.1.22 FSCA with ASD RBSR. **C,** Correlation of EC 4.2.1.22 HUMAnN with ASD RBSR. **D,** Correlation of EC 6.3.2.2 FSCA with ASD CARS. **E,** Correlation of EC 6.3.2.2 HUMAnN with ASD CARS. *RBSR: Repetitive Behavior Scale–Revised. CARS: Childhood Autism Rating Scale. *: pvalue < 0.05, **: pvalue < 0.01, ***: pvalue < 0.001*.

Other involved metabolic network of the 16 microbial functions were related to oxaloacetic acid, Acetyl CoA, Inosine, S-Adenosyl-L-homocysteine, pyrocatechol, and phenol (Supplementary Figure 7). The first two networks (Supplementary Figure 7A-B) were part of the *Krebs* Cycle and related to bioenergetic fluxes. Mitochondrial Dysfunction is being a new target for ASD treatments[73]. Specially, Oxaloacetic acid can stimulate neurogenesis and activate brain mitochondrial biogenesis[74]. Inosine (Supplementary Figure 7C) was thought to be a potential strategy of neurodegenerative diseases treatment[75, 76]. S-Adenosyl-L-homocysteine(Supplementary Figure 7D) were significantly increased in ASD[77] and this metabolite can be quickly linked to the glutamic acid-homocysteine metabolic network in Figure 6a as L-homocysteine is the precursor of S-Adenosyl-L-homocysteine. As for pyrocatechol (Supplementary Figure 7E), its derivate pyrocatechol sulfate was found significantly decreased in cerebrospinal fluid (CSF) of germ-free (GF) mice and might directly affect brain function and fear extinction learning[78]. 2-Naphthol (Supplementary Figure 7F) Levels were positively correlated with children’s allergic diseases[79]. Consistently, it has been reported that phenol sulfotransferase activities were decreased in ASD and associated with hyperserotonemia symptom[80].

For the 16 microbial functions apart from EC 4.2.1.22 and EC 6.3.2.2, we also observed stronger correlation between RFW microbial function and ASD symptoms than HUMAnN (Supplementary Figure 8). In short, through RFW, we proved that HUMAnN impaired the associations due to the distortion of microbial function information.

## Discussion

Current microbial functional profiling method exhibited severe information loss on both function detection and abundance calculation. To make complement for the loss, we developed a new framework, RFW, for microbial function inference on WMS, with aids of an annotated genome function reference database in this work. Previous reference-based functional profile inference methods, i.e., PICRUSt2[32, 34] and Tax4Fun2[81, 82] were designed for genus-level function inference based on16S rRNA gene sequences. Our work was the first species-level function inference method for WMS sequencing data.

We demonstrated the number of microbial functions reported by HUMAnN on WMS reduced sharply when relative abundance of taxon was lower than 5%-10%, which means if we want a complete microbial function information for 1% abundance taxa, a deep sequencing with 10x current sequencing depth would be required, and 100x for 0.1% abundance taxa, etc. This definitely would bring out a high experimental cost. Even for WMS on bacterial isolates, HUMAnN still theoretically failed to reported multiple microbial functions which had been registered in KEGG and UniProt. This loss might happen in library preparation or alignment search on the short DNA reads. The analysis on HUMAnN detection preference revealed several ongoing lost microbial pathways, involving in cholesterol and tryptophan metabolism, etc. Obviously, the information loss would harness our understanding on the potential functional capability of microbiome.

RFW was designed to predict microbial functional profile from WMS taxonomic profile using the genome-function reference database. Certainly, accuracy of RFW depends on accurate species identification and abundance estimation. Although this study cannot benchmark the accuracy of taxonomic profiling methods, we evaluated RFW abundance from Kraken2 and Metaphlan4, which were both commonly used in WMS taxonomic profiling. It was shown there was a highly consistency between RFW abundance from the two methods for co-detected microbial functions. Meanwhile, for either taxonomic profiling methods, RFW abundance revealed significantly more microbial functions than HUMAnN abundance. Researches has been devoted to improve taxon abundance estimation accuracy[83], we also expect to embrace more accurate taxonomic profiling methods in the future.

Benefited from completeness of RFW profile, the differential microbial function identification performance was greatly improved. Besides, we proposed a new hypothesis test, DFSCA-BC for absolute microbial functional abundance changes quantification and differential microbial function identification between groups. RFW and DFSCA-BC highlights several changed microbial functions related to colorectal cancer patients. RFW also supported associations construction between gut microbial functions and autism symptoms. An interesting point is that multiple studies could be found on explicating the role of corresponding Homo sapiens isoenzymes, while there are no studies to investigate the role of the highlighted enzymes from microbiome.

RFW provided support for functional profiling at the species and community levels. The genome-function reference database was built from function annotation on representative microbe genomes from GTDB database. Due to lacking of strain-level genomes, RFW could not generate functional profiles or distinguish functional variation at the strain level, which is the limitation of our approach.

In summary, RFW provides a comprehensive and complete species level scene of the complex microbial systems. The full functional information could be helpful in diverse downstream modeling tasks, i.e., differential microbial function identification, association analysis between microbial function and metabolites, etc. We believe that RFW will be a nice WMS functional profiling tool for renewing our knowledge on the host-microbe interactions.

## MATERIALS AND METHODS

### Taxon identification and HUMAnN analysis of WMS on bacterial isolates

Contigs were assembled by megahit v1.2.9[84], indexed by bowtie2 v2.3.5.1[85], binned by Metabat2 v2.12.1[86]. The generated MAGs was CheckM2 v1.0.1[87] and taxonomy assigned by GTDB-Tk v2.2.4[88]. The functional profiling of WMS on bacterial isolates was analyzed by HUMAnN 3.6.1[49](database: full_mapping v201901b, full_chocophlan v201901_v31, uniref90_annotated v201901b_full, MetaPhlAn v4.0.3 mpa_vJan21 CHOCOPhlAnSGB 202103).

### Taxonomic profiling and functional profiling of WMS on gut microbiome

WMS sequencing depth was accessed by FastQC v0.12.1[89]. Adapter and low quality reads were removed by Trimmomatic v0.39[90]. Contamination reads were removed by bowtie2 2.3.5.1[85] with H. sapiens GRCh38 + major SNVs. The Kraken2 based taxonomic profiling were performed by Kraken v2.1.2[47, 91] and braken v2.8[47, 92] with Kraken Standard plus protozoa & fungi 3/14/2023 Refseq indexes. The Metaphlan4 based taxonomic profiling were performed by MetaPhlAn v4.0.3 with mpa_vJan21 CHOCOPhlAnSGB_202103. Functional profiling of WMS was done by HUMAnN 3.6.1[49](database: full_mapping v201901b, full_chocophlan v201901_v31, uniref90_annotated v201901b_full, MetaPhlAn v4.0.3 mpa_vJan21 CHOCOPhlAnSGB 202103).

### RFW reference database building

With eggNOG-mapper (V2.1.4-2)[93, 94], we annotated protein-coding gene functions of representative genomes from 62,291 bacteria and 3,412 archaea species in GTDB (Genome Taxonomy Database, r207)[95]. The annotation database was based on eggNOG 5.0[96].

We also considered the 7302 genome-scale metabolic reconstruction (GSMR) models in AGORA2[97] as our second choice because GSMR models were also used as one alternative source of microbial activities in researches, e.g., MIMOSA2[98].

Microbial functions achieved from KEGG served as a third party one to validate the Genome-annotated or AGORA2 information sources. Potential functions of three common bacteria, *Escherichia coli UMN026* (Supplementary Figure 9a), *Bacteroides thetaiotaomicron VPI 5482* (Supplementary Figure 9b) and *Lactobacillus reuteri DSM 20016* (Supplementary Figure 9c), were mapped against each other by Venn plot from the above three sources. The encoded enzymes from the genome-annotation result were in agreement with that from KEGG at 90 percent. In contrast, AGORA2 missed much more microbial enzymes compared with the genome-annotation result or KEGG. For example, *Escherichia coli UMN026* could function in over 1101 enzymes from prediction of KEGG, in which 1022 enzymes (92.8%) were also annotated from the genome; AGORA2 only included 515 enzymes (46.8%) from KEGG. Analysis on the remained two bacteria supported the similar conclusion. We then calculated the common EC ratio for 6964 species-level AGORA2 models and revealed that AGORA2 only covered approximately 40% genome functions (Supplementary Figure 9d).

Conclusively, we have more confidence in the genome-annotation result rather than the AGORA2 model in microbial function profiling given its consistency with KEGG. Accordingly, the genome-annotation on GTDB database was used as the default RFW reference database.

### The RFW algorithm implementation

RFW was implemented as open source. The basic algorithm was coded in python and the genome-function reference database was managed by SQLite3. To map species identified from taxonomic profiling methods, RFW was designed to support both NCBI tax id and GTDB Taxonomy identifier. NCBI tax id mapping was used for Kraken2 based profiling. For Metaphlan4 derived taxonomic profiling, we used command sgb_to_gtdb_profile for transfering Metaphlan SGB to GTDB Taxonomy identifier. RFW yields mapped and no-mapped taxa information at this process.

FA and FSCA were inferred for mapped microbial species, with function encoding and function counts information extracted from the reference database. RFW support KO, EC and COG for describing microbial function. The reference database also stored 3445 Metacyc pathways which were used for microbial pathway coverage and abundance calculation. Besides, RFW also yields the sub-community composition for each microbial function and pathway. The composition could be used to determine the taxon contribution in the sub-community.

## Data & Code availability

All data for the article and the supplemental figures analysis and visualization are available on figshare https://www.doi.org/10.6084/m9.figshare.24936453. The microbial genome function reference database is available on figshare https://doi.org/10.6084/m9.figshare.24541876. RFW and DFSCA-BC are available at https://github.com/Xingyinliu-Lab/RFW. Code used for data analysis in this word are available on figshare https://www.doi.org/10.6084/m9.figshare.24936453.

## Acknowledgements

This work was supported by the National Key Research and Development Program, 2022YFA1303900; the National Natural Science Foundation of China (NSFC) grant 82172288; the Key society development project of Jiangsu Province, BE2021721.

## Author Contributions

X.L and K.M. conceived and designed this study. K.M developed the methodology, constructed the software, performed analysis, generated figures and wrote the manuscript. XL revised manuscript. All authors read, checked and approved the final manuscript.

## Competing interests

The authors declare no competing interests.

## Ethics statement

Not applicable.

## Supplementary Data

**Supplementary Figure 1.**
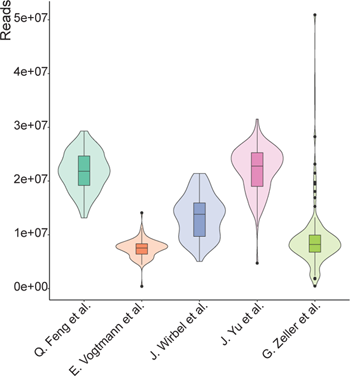
Sequencing reads count of WMS from different cohorts.

**Supplementary Figure 2.**
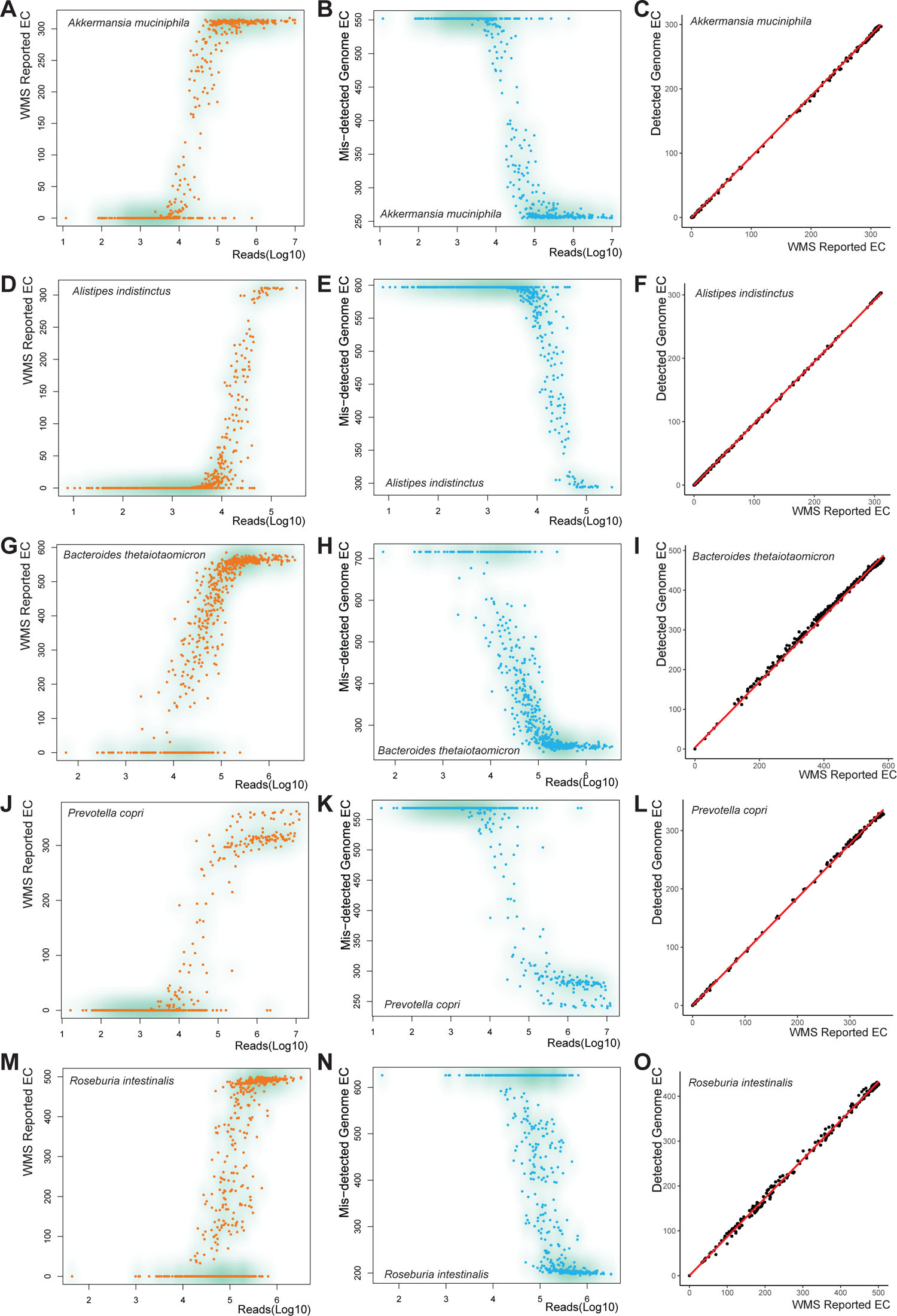
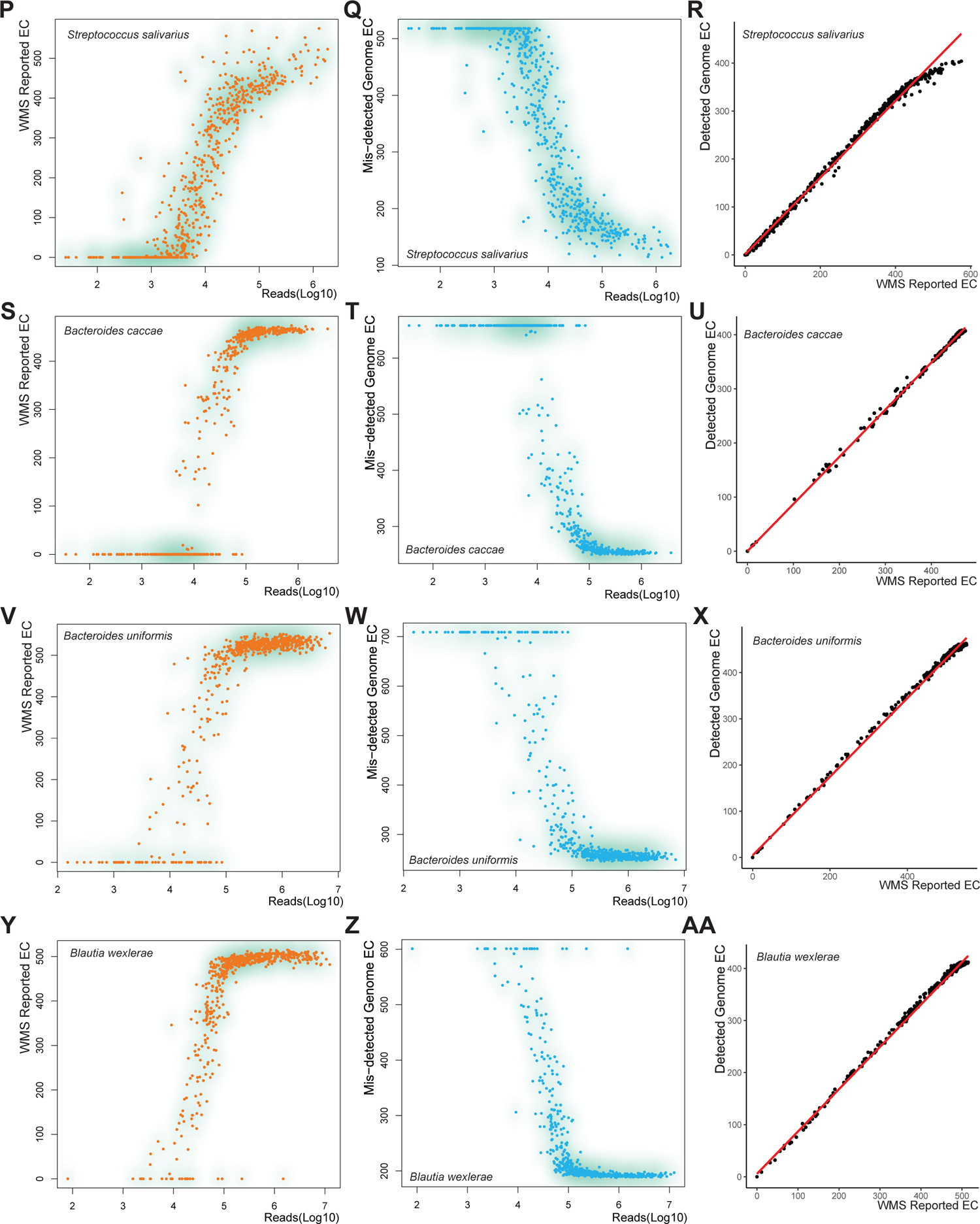
Taxon-level function loss in HUMAnN was connected to sequencing depth (assessed by Kraken2). HUMAnN reported enzymes of *Akkermansia muciniphila* (**A**), *Alistipes indistinctus* (**D**), *Bacteroides thetaiotaomicron* (**G**), *Prevotella copri* (**J**), *Roseburia intestinalis* (**M**), *Streptococcus salivarius* (**P**), *Bacteroides caccae* (**S**), *Bacteroides uniformis* (**V**), *Blautia wexlerae* (**Y**) from WMS on gut microbiome degraded sharply with shallow sequencing depth (by reads). Missed genome-annotated enzymes increased with shallow sequencing depth of *Akkermansia muciniphila* (**B**), *Alistipes indistinctus* (**E**), *Bacteroides thetaiotaomicron* (**H**), *Prevotella copri* (**K**), *Roseburia intestinalis* (**N**), *Streptococcus salivarius* (**Q**), *Bacteroides caccae* (**T**), *Bacteroides uniformis* (**W**), *Blautia wexlerae* (**Z**) from WMS on gut microbiome. Taxon-level enzymes reported by HUMAnN had a positive linear correlation with the covered genome annotated enzymes for *Akkermansia muciniphila* (**C**), *Alistipes indistinctus* (**F**), *Bacteroides thetaiotaomicron* (**I**), *Prevotella copri* (**L**), *Roseburia intestinalis* (**O**), *Streptococcus salivarius* (**R**), *Bacteroides caccae* (**U**), *Bacteroides uniformis* (**X**), *Blautia wexlerae* (**AA**).

**Supplementary Figure 3.**
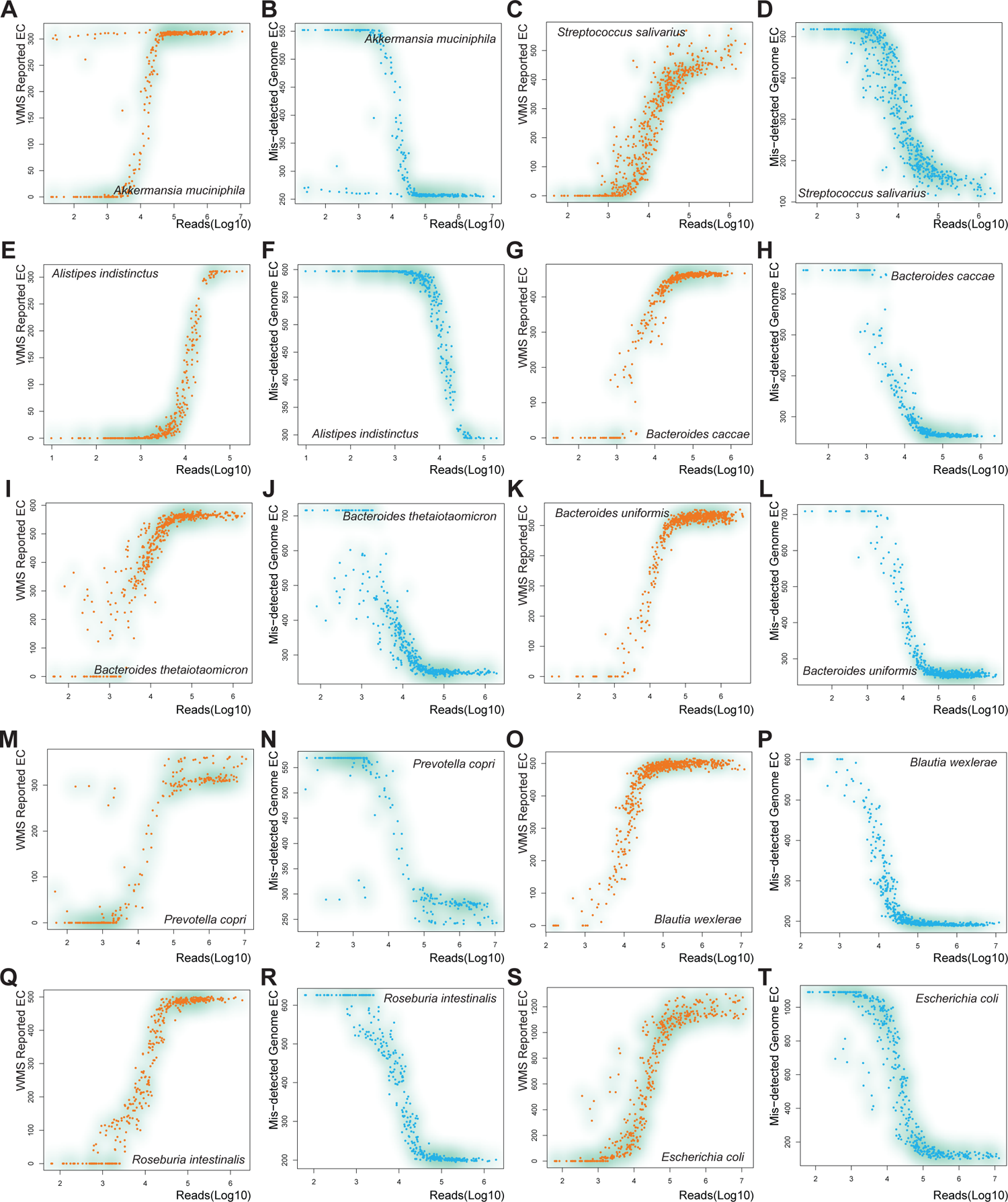
Taxon-level function loss in HUMAnN was connected to sequencing depth (assessed by Metaphlan4). HUMAnN reported enzymes of *Akkermansia muciniphila* (**A**), *Alistipes indistinctus* (**E**), *Bacteroides thetaiotaomicron* (**I**), *Prevotella copri* (**M**), *Roseburia intestinalis* (**Q**), *Streptococcus salivarius* (**C**), *Bacteroides caccae* (**G**), *Bacteroides uniformis* (**K**), *Blautia wexlerae* (**O**), *Escherichia coli* (**S**) from WMS on gut microbiome degraded sharply with shallow sequencing depth (by reads). Missed genome-annotated enzymes increased with shallow sequencing depth of *Akkermansia muciniphila* (**B**), *Alistipes indistinctus* (**F**), *Bacteroides thetaiotaomicron* (**J**), *Prevotella copri* (**N**), *Roseburia intestinalis* (**R**), *Streptococcus salivarius* (**D**), *Bacteroides caccae* (**H**), *Bacteroides uniformis* (**L**), *Blautia wexlerae* (**P**), *Escherichia coli* (**T**) from WMS on gut microbiome.

**Supplementary Figure 4.**
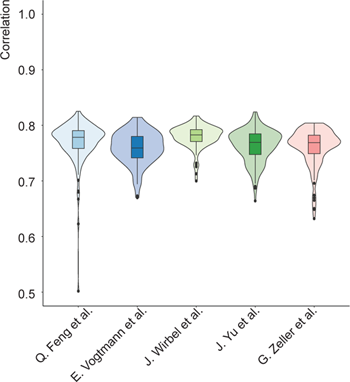
RFW-FA with taxonomic profiling from Metaphlan4 had a strong positive Pearson correlation with HUMAnN reported functional abundance.

**Supplementary Figure 5.**
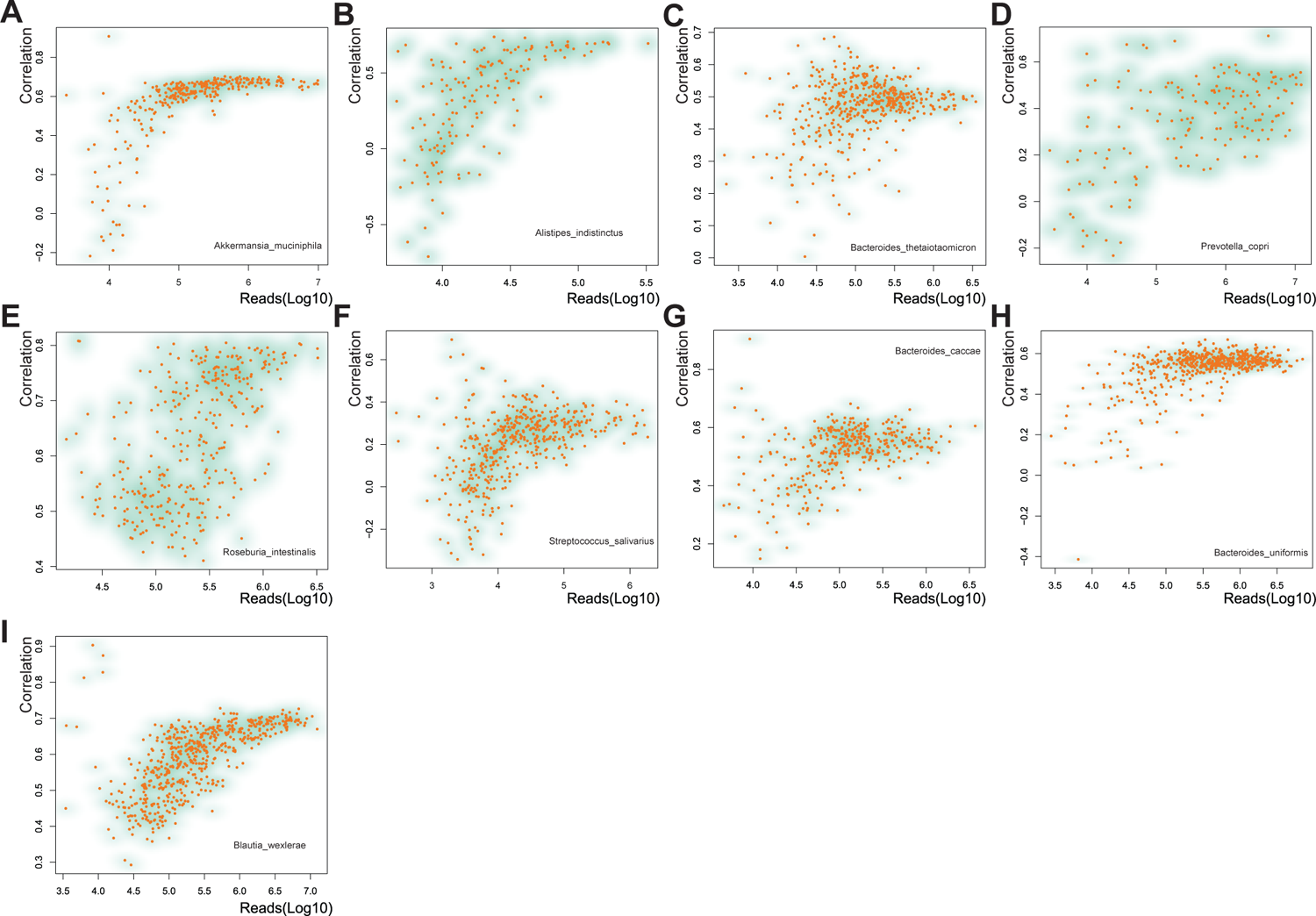
The sequence depth affected correlations between taxon-level RFW-FA (using taxonomic profile from Kraken2) and HUMAnN. **A-I,** The sequence depth affected correlations between functional abundance of *Akkermansia muciniphila* (**A**), *Alistipes indistinctus* (**B**), *Bacteroides thetaiotaomicron* (**C**), *Prevotella copri* (**D**), *Roseburia intestinalis* (**E**), *Streptococcus salivarius* (**F**), *Bacteroides caccae* (**G**), *Bacteroides uniformis* (**H**), *Blautia wexlerae* (**I**) from RFW and from HUMAnN.

**Supplementary Figure 6.**
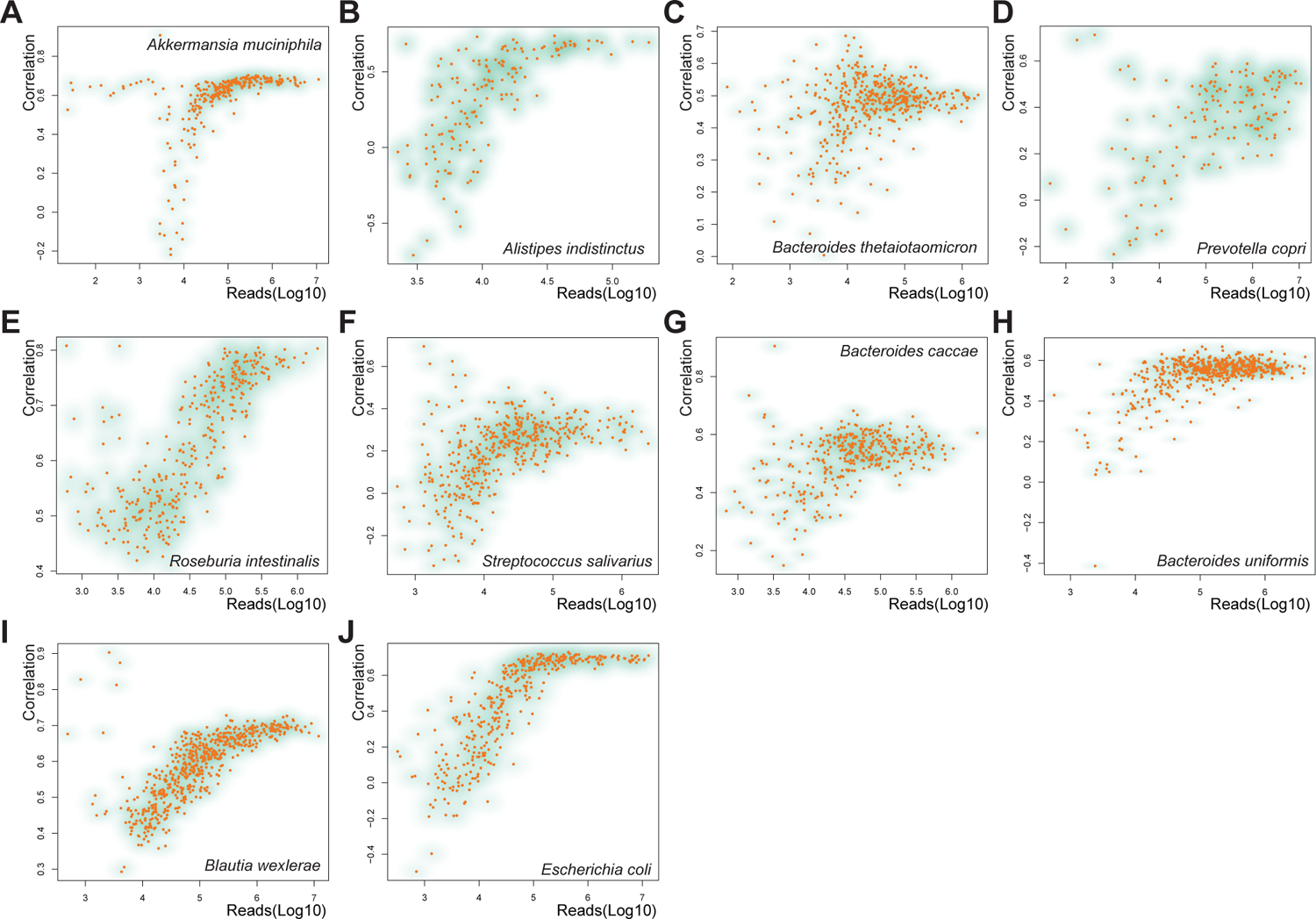
The sequence depth affected correlations between taxon-level RFW-FA (using taxonomic profile from Metaphlan4) and HUMAnN. **A-J,** The sequence depth affected correlations between functional abundance of *Akkermansia muciniphila* (**A**), *Alistipes indistinctus* (**B**), *Bacteroides thetaiotaomicron* (**C**), *Prevotella copri* (**D**), *Roseburia intestinalis* (**E**), *Streptococcus salivarius* (**F**), *Bacteroides caccae* (**G**), *Bacteroides uniformis* (**H**), *Blautia wexlerae* (**I**), *Escherichia coli* (**J**) from RFW and from HUMAnN.

**Supplementary Figure 7.**
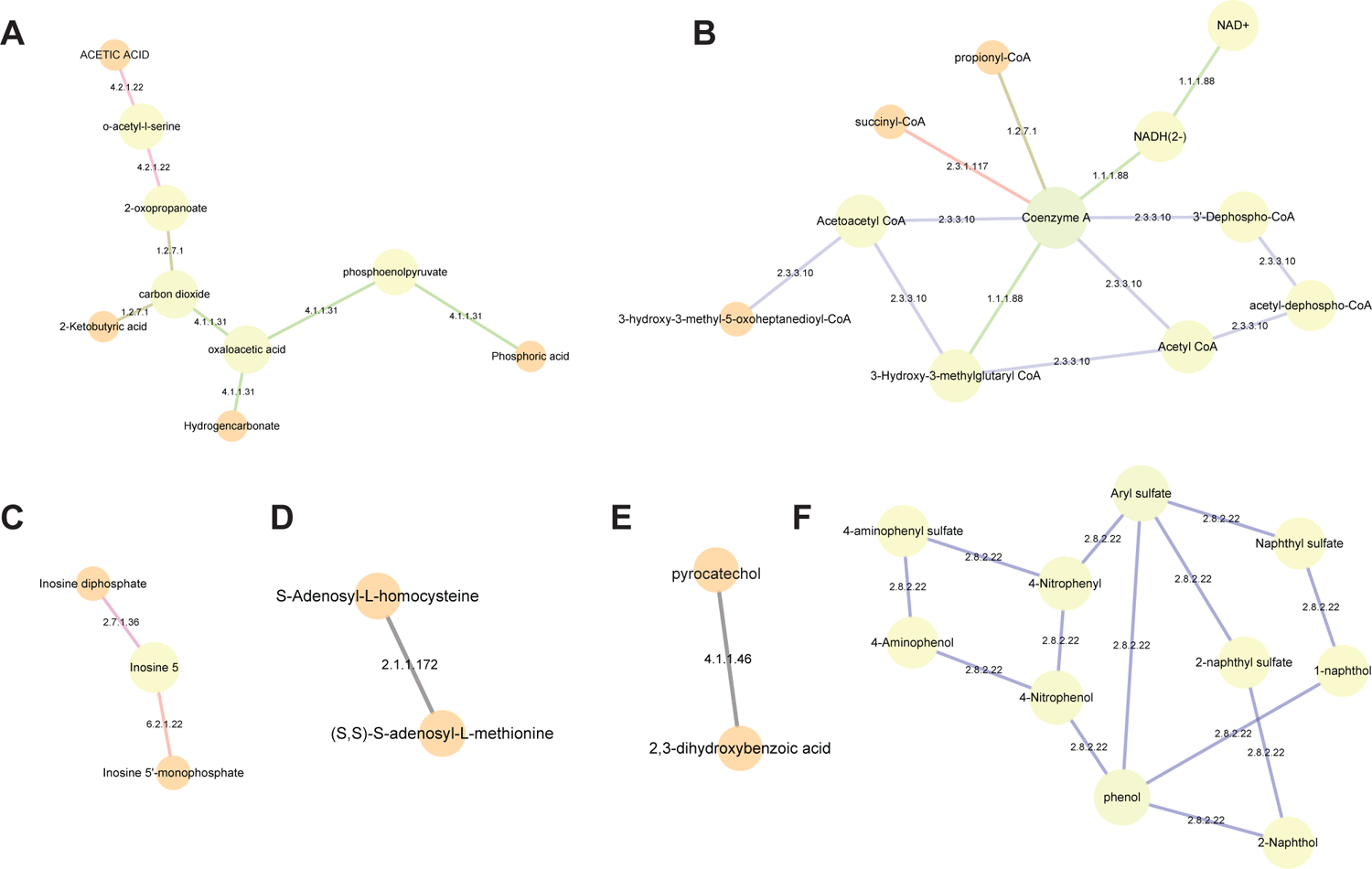
metabolic network of ASD symptom associated microbial functions. **A,** EC 1.2.7.1, EC 4.1.1.31 and EC 4.2.1.22 involved metabolic network. **B,** EC 1.1.1.88, EC 1.2.7.1, EC 2.3.1.117 and EC 2.3.3.10 involved metabolic network. **C,** EC 2.7.1.36 and EC 6.2.1.22 involved metabolic network. **D,** EC 2.1.1.172 involved metabolic network. **E,** EC 4.1.1.46 involved metabolic network. **F,** EC 2.8.2.22 involved metabolic network.

**Supplementary Figure 8.**
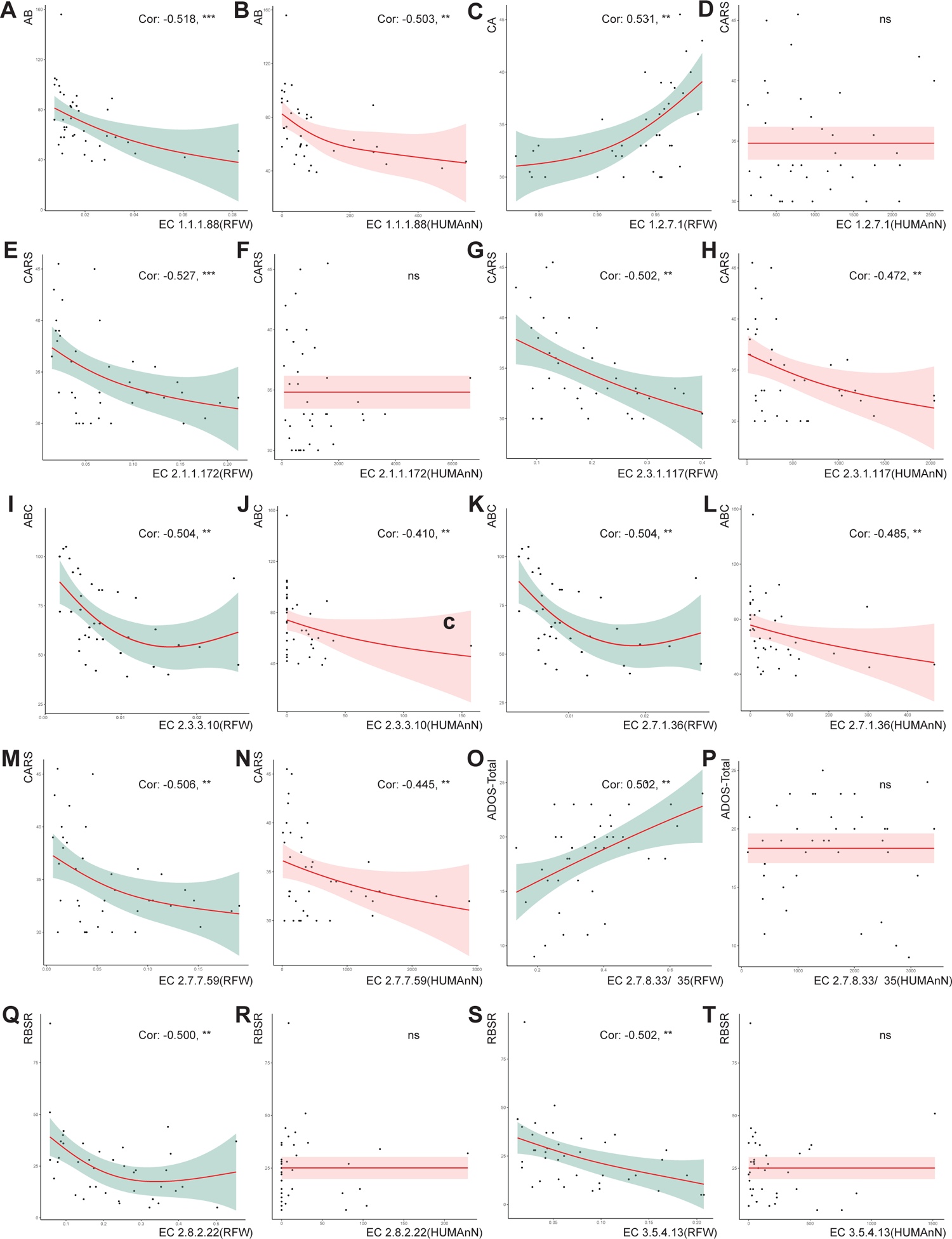
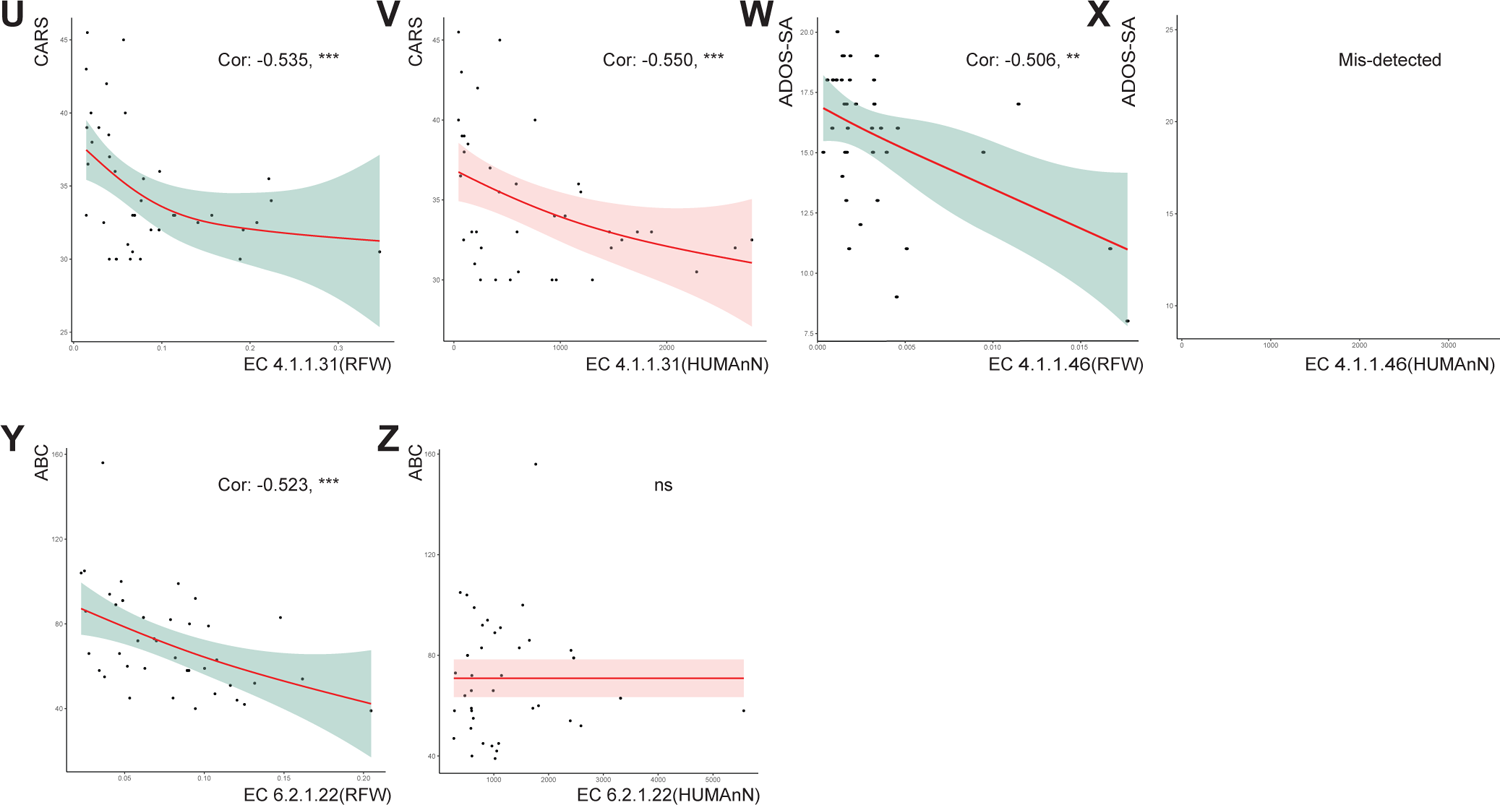
association between microbial functions and ASD symptom. **A,** Correlation of EC 1.1.1.88 FSCA with ASD ABC. **B,** Correlation of EC 1.1.1.88 HUMAnN with ASD ABC. **C,** Correlation of EC 1.2.7.1 FSCA with ASD CARS. **D,** Correlation of EC 1.2.7.1 HUMAnN with ASD CARS. **E,** Correlation of EC 2.1.1.172 FSCA with ASD CARS. **F,** Correlation of EC 2.1.1.172 HUMAnN with ASD CARS. **G,** Correlation of EC 2.3.1.117 FSCA with ASD CARS. **H,** Correlation of EC 2.3.1.117 HUMAnN with ASD CARS. **I,** Correlation of EC 2.3.3.10 FSCA with ASD ABC. **J,** Correlation of EC 2.3.3.10 HUMAnN with ASD ABC. **K,** Correlation of EC 2.7.1.36 FSCA with ASD ABC. **L,** Correlation of EC 2.7.1.36 HUMAnN with ASD ABC. **M,** Correlation of EC 2.7.7.59 FSCA with ASD CARS. **N,** Correlation of EC 2.7.7.59 HUMAnN with ASD CARS. **O,** Correlation of EC 2.7.8.33/35 FSCA with ASD ADOS-Total. **P,** Correlation of EC 2.7.8.33/35 HUMAnN with ASD ADOS-Total. **Q,** Correlation of EC 2.8.2.22 FSCA with ASD RBSR. **R,** Correlation of EC 2.8.2.22 HUMAnN with ASD RBSR. **S,** Correlation of EC 3.5.4.13 FSCA with ASD RBSR. **T,** Correlation of EC 3.5.4.13 HUMAnN with ASD RBSR. **U,** Correlation of EC 4.1.1.31 FSCA with ASD CARS. **V,** Correlation of EC 4.1.1.31 HUMAnN with ASD CARS. **W,** Correlation of EC 4.1.1.46 FSCA with ASD ADOS-SA. **X,** EC 4.1.1.46 were not detected in HUMAnN. **Y,** Correlation of EC 6.2.1.22 FSCA with ASD ABC. **Z,** Correlation of EC 6.2.1.22 HUMAnN with ASD ABC. *RBSR: Repetitive Behavior Scale–Revised. CARS: Childhood Autism Rating Scale. ABC: Autism Behavior Checklist. ADOS: Autism Diagnostic Observation Schedule. *: pvalue < 0.05, **: pvalue < 0.01, ***: pvalue < 0.001*.

**Supplementary Figure 9.**
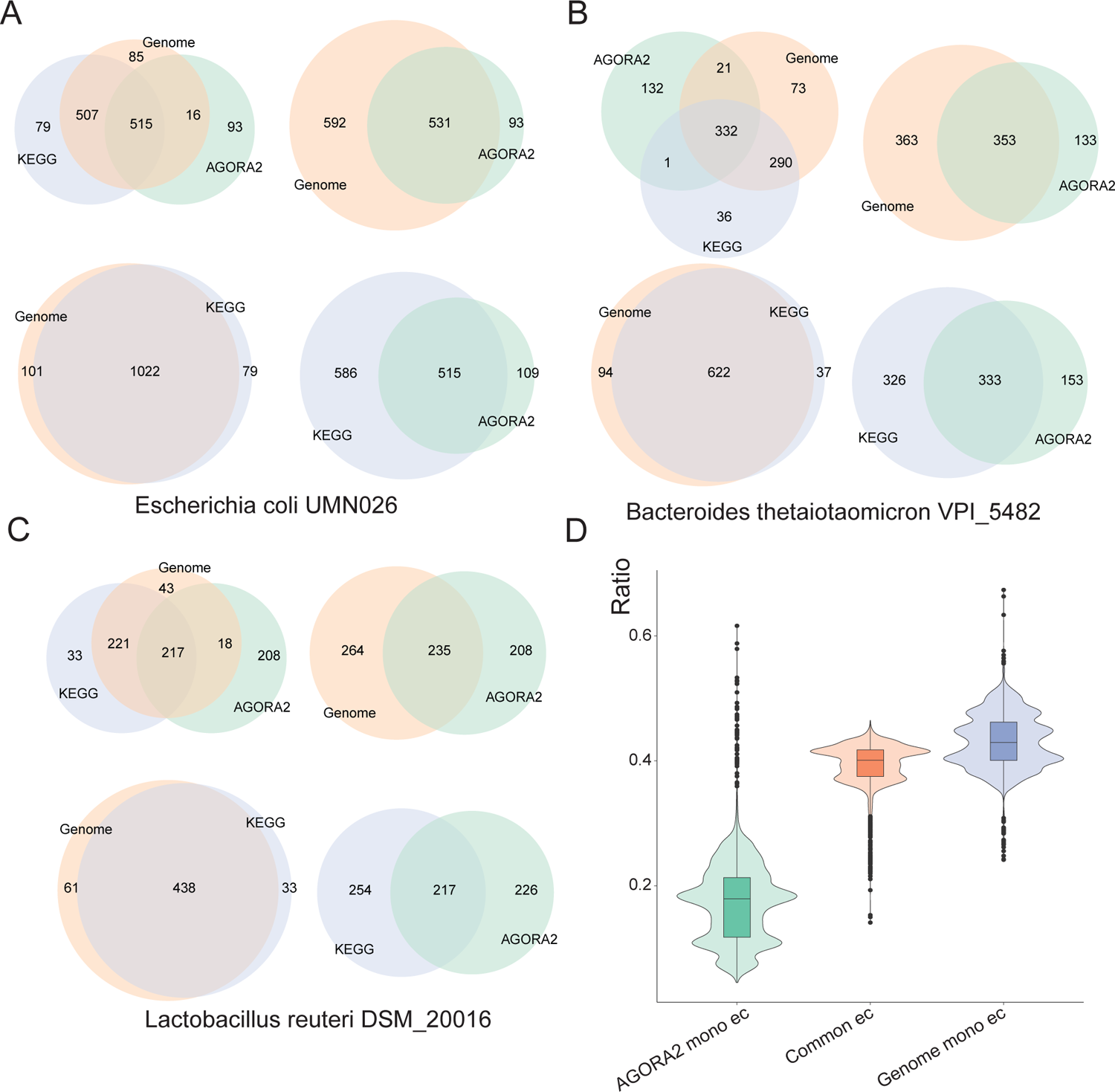
Genome-function reference database validation. **A-C,** *Venn plots of encoded 4th Level EC enzymes which were extracted from KEGG, genome-annotation and AGORA2 for Escherichia coli UMN026* (**A**), *Bacteroides thetaiotaomicron VPI 5482* (**B**), *Lactobacillus reuteri DSM 20016* (**C**). **D,** AGORA2 covered approximately 40% genome functions. *AGORA2 mono EC: ratio between counts of enzymes monopolized in AGORA2 and counts of all enzymes. Common EC: ratio between counts of common enzymes shared by both AGORA2 & genome-annotation and counts of all enzymes. Genome mono EC: ratio between counts of enzymes monopolized in genome-annotation and counts of all enzymes. The counts of all enzymes were calculated as number of enzymes collected from AGORA2 and genome-annotation with duplicates filtered*.

### Supplementary information 1: Proof on formula 1

Without loss of generality, suppose the sub-community ℒ for encoding funcition *f* were composed with two taxa, *T*, *T*, i.e., ℒ = {*Taxon*|φ^*f*^ = 1} = {*T*, *T* }. We have,

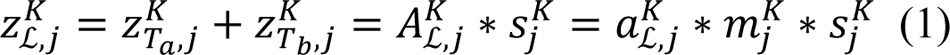

Where *A*^*K*^_ℒ, j_ represents the total number taxa in sub-community ℒ contained in Group K, sample j. *a*^*K*^_ℒ, j_ represents the absolute abundance of taxa in sub-community ℒ in Group K, sample j. *m*^*K*^_j_ represents the sampling mass of Group K, sample j. *S*^*K*^_j_ is the sampling fraction of Group K, sample j. *Z*^*K*^_ℒ, j_, *Z*^*K*^_Ta, j_, and *Z*^*K*^_Tb, j_ represents sequencing read counts of *Taxa*_*i*_ in Group K, sample j. K denotes the group index. For simplicity of exposition, K=T, C, namely the C (Control) group and the T (Treatment) Group.

Then the relative abundance of sub-community ℒ, i.e. the *FSCA*^*K*^_f,j_ for function *f* could be calculated as:

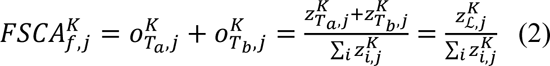

Hence,

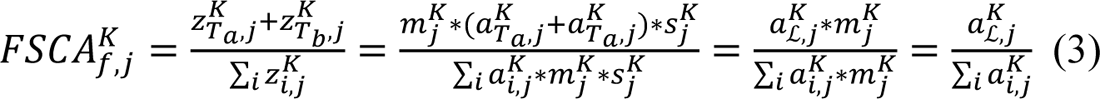

i.e.,

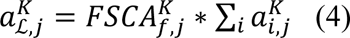

where ∑_*i*_ *a*^*K*^ is the total absolute abundance of all taxa. Denote Ψ_j_^*K*^ = ∑_*i*_ *a*^*K*^,

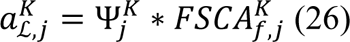

we have Thus, the absolute abundance change Δ_*f*_ of microbial function *f* is

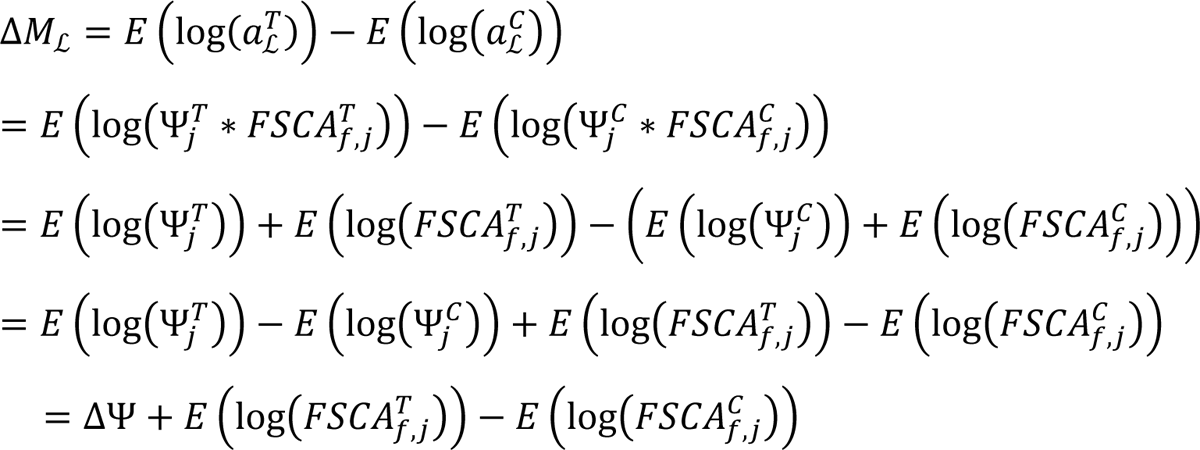

where ΔΨ = *E* (log(Ψ_j_^*T*^)) − *E* (log(Ψ_j_^*C*^)) is the log fold change of total microbial absolute abundance of all taxa between groups. Q.E.D.

**Supplementary Table 1.** Genome information on 15 bacterial isolate WMS.

| Reference reported |  | Mapped GTDB genome information |  |  |  |
| --- | --- | --- | --- | --- | --- |
| Bacterial strains<sup>[1]</sup> | WMS sequence<sup>[2]</sup> | Refseq assembly id | Phylum | Species | NCBI Species Taxid |
| *Escherichia coli* Mt1B1 | SRR9696643 | GCF_003697165.2 | Proteobacteria | *Escherichia coli* | 562 |
| *Irregularicoccus caecimuris* MD308 | SRR9696644 | GCA_010206225.2 | Firmicutes A | MD308 sp010206225 | 1235799 |
| *Anaerotruncus colihominis* JM4-15 | SRR9696645 | GCF_010206195.1 | Firmicutes A | *Anaerotruncus colihominis* A | 169435 |
| *Enterocloster clostridioformis* YL32 | SRR9696646 | GCF_900113155.1 | Firmicutes A | *Enterocloster clostridioformis* | 1531 |
| *Clostridium cocleatum* I50 | SRR9696647 | GCF_900102365.1 | Firmicutes | *Erysipelatoclostridium cocleatum* | 69824 |
| *Bacteroides acidifaciens* MD185 | SRR9696648 | GCA_000613385.1 | Bacteroidota | *Bacteroides acidifaciens* | 85831 |
| *Bacteroides caecimuris* MD237 | SRR9696649 | GCF_001688725.2 | Bacteroidota | *Bacteroides caecimuris* | 1796613 |
| *Clostridium* sp. MD294 | SRR9696650 | GCF_000364165.1 | Firmicutes A | *Coprocola* sp000364165 | 97138 |
| *Clostridium* sp. MD300 | SRR9696651 | GCF_003885045.1 | Firmicutes A | *Schaedlerella arabinosiphila* | 2044587 |
| *Subtilibacillum caecimuris* MD335 | SRR9696652 | GCA_000403335.2 | Firmicutes A | COE1 sp000403335 | 1235793 |
| *Lactobacillus johnsonii* MD006 | SRR9696653 | GCF_000159355.1 | Firmicutes | *Lactobacillus johnsonii* | 33959 |
| *Ligilactobacillus murinus* MD040 | SRR9696654 | GCF_001591685.1 | Firmicutes | *Ligilactobacillus murinus* | 1622 |
| *Limosilactobacillus reuteri* MD207 | SRR9696655 | GCA_003072625.1 | Firmicutes | *Limosilactobacillus reuteri* D | 1598 |
| *Parabacteroides goldsteinii* MD072 | SRR9696656 | GCF_000969835.1 | Bacteroidota | *Parabacteroides goldsteinii* | 328812 |
| *Longibacillum caecimuris* MD329 | SRR9696657 | GCA_010206265.1 | Firmicutes A | CAG-41 sp010206265 | 2592090 |

- 2 - 3 [1] Darnaud M, De Vadder F, Bogeat P, et al. A standardized gnotobiotic mouse model harboring - 4 a minimal 15-member mouse gut microbiota recapitulates SOPF/SPF phenotypes[J]. *Nature* - 5 *Communications*, 2021, 12(1): 6686. - 6 [2] Elie C, Mathieu A, Saliou A, et al. Draft Genome Sequences of 15 Bacterial Species - 7 Constituting the Stable Defined Intestinal Microbiota of the GM15 Gnotobiotic Mouse - 8 Model[J]. *Microbiol Resour Announc*, 2020, 9(35).

**Supplementary Table 2.**
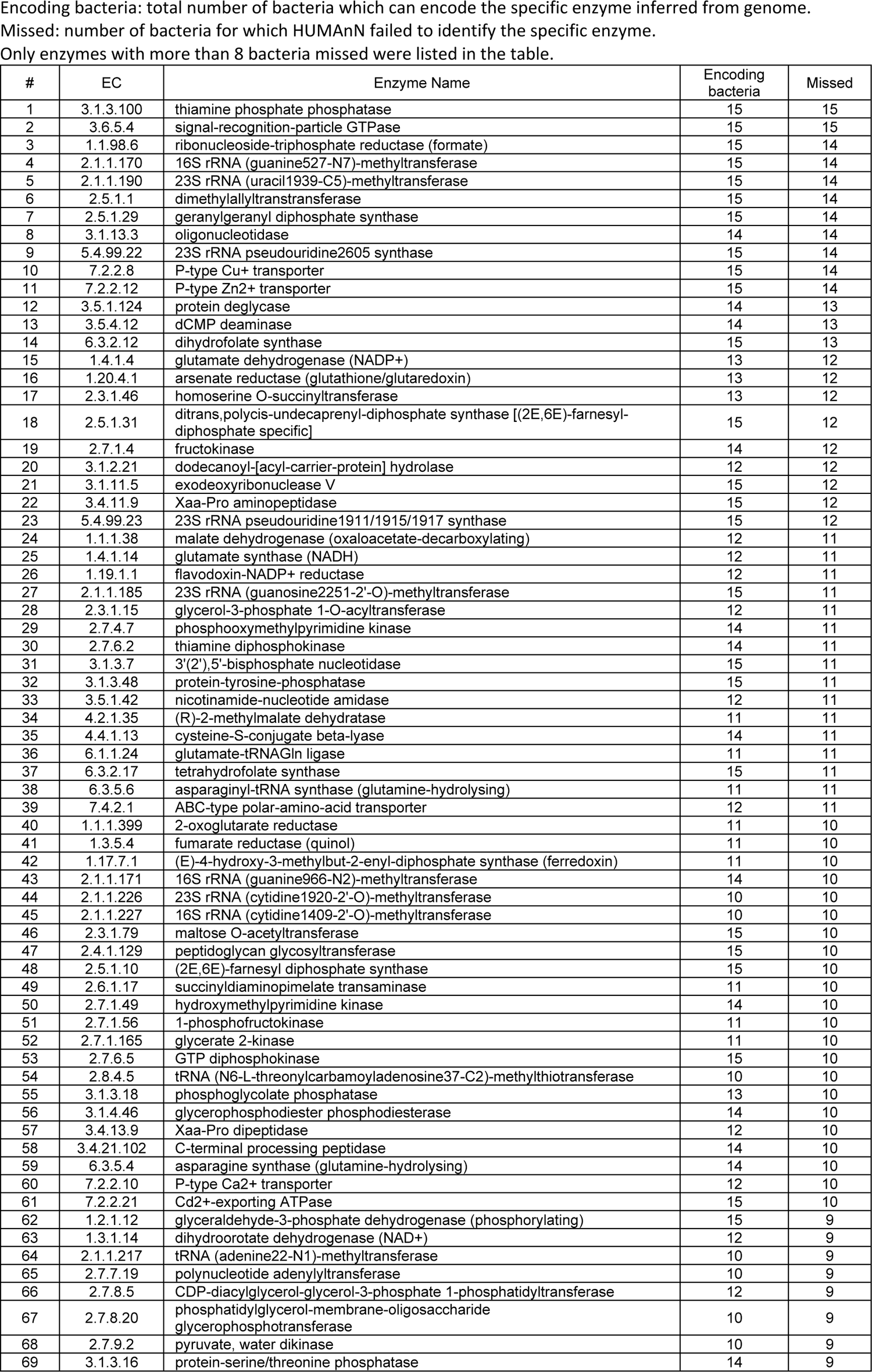
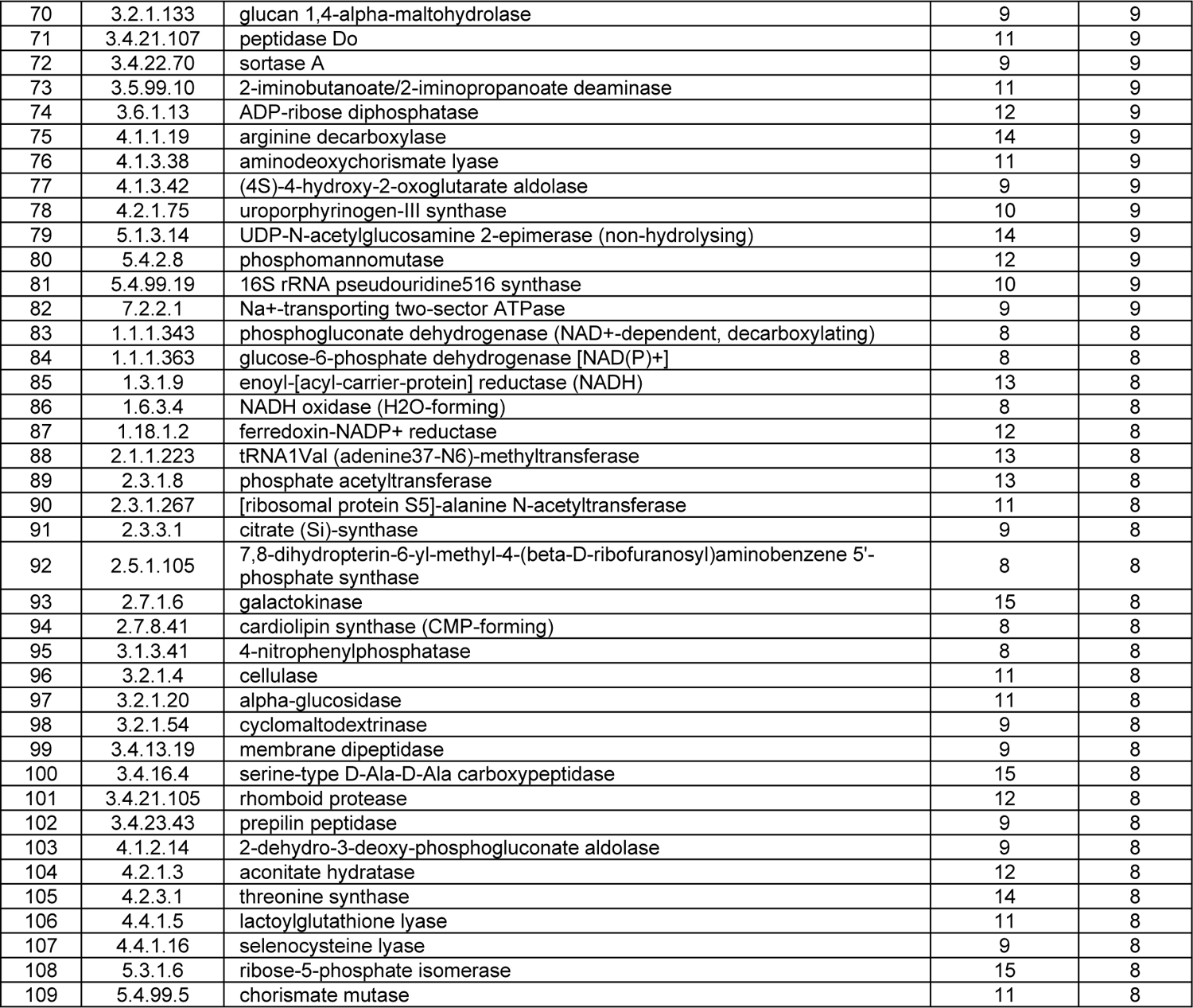
HUMAnN mis-detected bacteria functions from WMS on cultured isolates.

**Supplementary Table 3.** Evidence from KEGG and UniProt on bacterial enzyme encoding.

| Bacterial strains | EC 3.1.3.100<br>thiamine phosphate phosphatase |  | EC 3.6.5.4<br>signal-recognition-particle GTPase |  |
| --- | --- | --- | --- | --- |
|  | Gene | Evidence | Gene | Evidence |
| Escherichia coli<br>Mt1B1 | rsgA | • https://www.kegg.jp/entry/eco:b4161<br>• https://www.uniprot.org/uniprotkb/C3SG82/entry | ffh | • https://www.kegg.jp/entry/eco:b2610<br>• https://www.uniprot.org/uniprotkb/A0A376NUY1/entry |
| Irregularicoccus<br>caecimuris MD308 |  | • https://www.uniprot.org/uniprotkb/R9MBK4/entry |  | • https://www.uniprot.org/uniprotkb/R9MDY0/entry |
| Anaerotruncus<br>colihominis JM4-15 |  | • https://www.kegg.jp/entry/acol:K5I23_09750<br>• https://www.uniprot.org/uniprotkb/A0A845QL05/entry |  | • https://www.kegg.jp/entry/acol:K5I23_03885<br>• https://www.uniprot.org/uniprotkb/A0A845QJW0/entry |
| Enterocloster<br>clostridioformis YL32 |  | • https://www.uniprot.org/uniprotkb/A0A8B2X8Z4/entry |  | • https://www.uniprot.org/uniprotkb/A0A174C1H6/entry |
| Clostridium<br>cocleatum I50 |  | • https://www.uniprot.org/uniprotkb/A0A110EQF3/entry |  | • https://www.uniprot.org/uniprotkb/A0A829ZBN1/entry |
| Bacteroides<br>acidifaciens MD185 |  | • https://www.uniprot.org/uniprotkb/A0A4S2AM92/entry |  | • https://www.uniprot.org/uniprotkb/A0A4S2ADT6/entry |
| Bacteroides<br>caecimuris MD237 |  | • https://www.kegg.jp/entry/bcae:A4V03_14240<br>• https://www.uniprot.org/uniprotkb/A0A1C7H209/entry |  | • https://www.kegg.jp/entry/bcae:A4V03_12080<br>• https://www.uniprot.org/uniprotkb/A0A1C7H0N1/entry |
| Clostridium sp.<br>MD294 |  | • https://www.uniprot.org/uniprotkb/N1ZIW9/entry |  | • https://www.uniprot.org/uniprotkb/N1ZMB7/entry |
| Clostridium sp.<br>MD300 |  | • https://www.uniprot.org/uniprotkb/A0A426DDS1/entry |  | • https://www.uniprot.org/uniprotkb/A0A426DF19/entry |
| Subtilibacillum<br>caecimuris MD335 |  | • https://www.uniprot.org/uniprotkb/A0A7X5L271/entry |  | • https://www.uniprot.org/uniprotkb/A0A7X5L4L3/entry |
| Lactobacillus<br>johnsonii MD006 |  | • https://www.kegg.jp/entry/ljf:FI9785_729<br>• https://www.uniprot.org/uniprotkb/A0A6B9HZD2/entry |  | • https://www.kegg.jp/entry/ljf:FI9785_750<br>• https://www.uniprot.org/uniprotkb/A0A267LVS0/entry |
| Ligilactobacillus<br>murinus MD040 |  | • https://www.kegg.jp/entry/lmur:CPS94_04250<br>• https://www.uniprot.org/uniprotkb/A0A837ISI1/entry |  | • https://www.kegg.jp/entry/lmur:CPS94_04180<br>• https://www.uniprot.org/uniprotkb/A0A837ISY6/entry |
| Limosilactobacillus<br>reuteri MD207 |  | • https://www.kegg.jp/entry/lre:Lreu_1168<br>• https://www.uniprot.org/uniprotkb/A0A347T5S7/entry |  | • https://www.kegg.jp/entry/lre:Lreu_1155<br>• https://www.uniprot.org/uniprotkb/A0A143Q353/entry |
| Parabacteroides<br>goldsteinii MD072 |  | • https://www.kegg.jp/entry/pgol:K6V26_07690<br>• https://www.uniprot.org/uniprotkb/A0A6G1Z7N6/entry |  | • https://www.kegg.jp/entry/pgol:K6V26_00995<br>• https://www.uniprot.org/uniprotkb/A0A0J6CR05/entry |
| Longibacillum<br>caecimuris MD329 |  | • https://www.uniprot.org/uniprotkb/A0A7X5KRX6/entry |  | • https://www.uniprot.org/uniprotkb/A0A7X5KQA0/entry |

**Supplementary Table 4.** the most involved 20 pathways with HUMAnN mis-detected enzymes (assessed by Kraken2)

| # | Pathway id | Pathway Name | Involved EC | WMS missed EC |
| --- | --- | --- | --- | --- |
| 1 | PWY-6954 | superpathway of aromatic compound degradation via 2-hydroxypentadienoate | 36 | 32 |
| 2 | PWY-2504 | superpathway of aromatic compound degradation via 3-oxoadipate | 32 | 29 |
| 3 | PWY-6146 | Methanobacterium thermoautotrophicum biosynthetic metabolism | 51 | 28 |
| 4 | PWY-5327 | superpathway of L-lysine degradation | 37 | 24 |
| 5 | PWY-5183 | superpathway of aerobic toluene degradation | 26 | 23 |
| 6 | PWY-5257 | superpathway of pentose and pentitol degradation | 35 | 23 |
| 7 | ALL-CHORISMATE-PWY | superpathway of chorismate metabolism | 52 | 20 |
| 8 | ERGOSTEROL-SYN-PWY | superpathway of ergosterol biosynthesis I | 23 | 17 |
| 9 | PWY-6957 | mandelate degradation to acetyl-CoA | 18 | 17 |
| 10 | PWY66-5 | superpathway of cholesterol biosynthesis | 22 | 16 |
| 11 | PWY-6947 | superpathway of cholesterol degradation II (cholesterol dehydrogenase) | 17 | 16 |
| 12 | PWY-6928 | superpathway of cholesterol degradation I (cholesterol oxidase) | 17 | 16 |
| 13 | PWY-6830 | superpathway of methanogenesis | 19 | 16 |
| 14 | PWY-6937 | superpathway of testosterone and androsterone degradation | 19 | 15 |
| 15 | BENZCOA-PWY | anaerobic aromatic compound degradation (Thauera aromatica) | 20 | 14 |
| 16 | PWY-7024 | superpathway of the 3-hydroxypropanoate cycle | 17 | 13 |
| 17 | PWY-6505 | L-tryptophan degradation XII (Geobacillus) | 13 | 13 |
| 18 | PWY-5529 | superpathway of bacteriochlorophyll a biosynthesis | 16 | 13 |
| 19 | PWY-5655 | L-tryptophan degradation IX | 13 | 13 |
| 20 | PWY-8397 | arachidonate metabolites biosynthesis | 21 | 13 |

**Supplementary Table 5.** the most involved 20 pathways with HUMAnN mis-detected enzymes (assessed by Metaphlan4)

| # | Pathway id | Pathway Name | Involved EC | WMS missed EC |
| --- | --- | --- | --- | --- |
| 1 | PWY-6146 | Methanobacterium thermoautotrophicum biosynthetic metabolism | 51 | 28 |
| 2 | PWY-6954 | superpathway of aromatic compound degradation via 2-hydroxypentadienoate | 36 | 28 |
| 3 | PWY-2504 | superpathway of aromatic compound degradation via 3-oxoadipate | 32 | 28 |
| 4 | PWY-5183 | superpathway of aerobic toluene degradation | 26 | 22 |
| 5 | PWY-5257 | superpathway of pentose and pentitol degradation | 35 | 20 |
| 6 | PWY-5327 | superpathway of L-lysine degradation | 37 | 19 |
| 7 | ALL-CHORISMATE-PWY | superpathway of chorismate metabolism | 52 | 18 |
| 8 | PWY-6957 | mandelate degradation to acetyl-CoA | 18 | 16 |
| 9 | PWY-6947 | superpathway of cholesterol degradation II (cholesterol dehydrogenase) | 17 | 15 |
| 10 | PWY-6928 | superpathway of cholesterol degradation I (cholesterol oxidase) | 17 | 15 |
| 11 | PWY-6830 | superpathway of methanogenesis | 19 | 15 |
| 12 | PWY-6937 | superpathway of testosterone and androsterone degradation | 19 | 14 |
| 13 | PWY-6505 | L-tryptophan degradation XII (Geobacillus) | 13 | 13 |
| 14 | PWY-5655 | L-tryptophan degradation IX | 13 | 13 |
| 15 | PWY-6944 | androstenedione degradation I (aerobic) | 13 | 12 |
| 16 | PWY-5178 | toluene degradation IV (aerobic) (via catechol) | 13 | 12 |
| 17 | PWY-5507 | adenosylcobalamin biosynthesis I (anaerobic) | 26 | 11 |
| 18 | ERGOSTEROL-SYN-PWY | superpathway of ergosterol biosynthesis I | 23 | 11 |
| 19 | PWY-5529 | superpathway of bacteriochlorophyll a biosynthesis | 16 | 11 |
| 20 | TRYPTOPHAN-DEGRADATION-1 | L-tryptophan degradation III (eukaryotic) | 14 | 11 |

**Supplementary Table 6.** enzyme detection in PWY-5655 L-tryptophan degradation IX.

| EC | Positive samples in HUMAnN | Assessed by Metaphlan4 |  |  | Assessed by Kraken2 |  |  |
| --- | --- | --- | --- | --- | --- | --- | --- |
|  |  | Genome function Positive samples | True positive samples | True positive rate | Genome function Positive samples | True positive samples | True positive rate |
| 1.13.11.11 | 3 | 32 | 3 | 9.38% | 589 | 3 | 0.51% |
| 1.13.11.52 | 0 | 9 | 0 | 0.00% | 587 | 0 | 0.00% |
| 1.13.11.6 | 42 | 106 | 11 | 10.38% | 589 | 42 | 7.13% |
| 1.14.13.9 | 13 | 90 | 2 | 2.22% | 589 | 13 | 2.21% |
| 1.2.1.10 | 534 | 589 | 534 | 90.66% | 589 | 534 | 90.66% |
| 1.2.1.32 | 89 | 588 | 89 | 15.14% | 589 | 89 | 15.11% |
| 3.5.1.9 | 571 | 589 | 571 | 96.94% | 589 | 571 | 96.94% |
| 3.5.99.5 | 342 | 16 | 6 | 37.50% | 589 | 342 | 58.06% |
| 3.7.1.3 | 8 | 375 | 8 | 2.13% | 589 | 8 | 1.36% |
| 4.1.1.45 | 0 | 427 | 0 | 0.00% | 589 | 0 | 0.00% |
| 4.1.1.77 | 142 | 539 | 140 | 25.97% | 589 | 142 | 24.11% |
| 4.1.3.39 | 579 | 588 | 578 | 98.30% | 589 | 579 | 98.30% |
| 4.2.1.80 | 293 | 546 | 285 | 52.20% | 589 | 293 | 49.75% |

**Supplementary Table 7.** microbial functions which were judged as differential ones between Health and CRC by Deseq2 on RFW-FA but not on HUMAnN in at least four cohorts.

| EC | EC name | Newly reported cohorts | RFW Positive Samples | HUMAnN Positive samples |
| --- | --- | --- | --- | --- |
| 2.4.1.152 | Lewis-negative α-3-fucosyltransferase | 5 | 589 | 0 |
| 2.4.1.65 | (Lea)-dependent (α-3/4)-fucosyltransferase | 5 | 589 | 0 |
| 1.4.1.11 | L-3,5-diaminohexanoate dehydrogenase | 4 | 589 | 271 |
| 2.1.1.137 | arsenite methyltransferase | 4 | 589 | 0 |
| 2.7.10.1 | receptor protein-tyrosine kinase | 4 | 589 | 343 |
| 3.4.24.13 | IgA-specific metalloendopeptidase | 4 | 589 | 336 |
| 4.2.1.120 | 4-hydroxybutanoyl-CoA dehydratase | 4 | 589 | 500 |
| 4.3.1.2 | methylaspartate ammonia-lyase | 4 | 589 | 206 |
| 5.4.3.3 | lysine 5,6-aminomutase | 4 | 589 | 158 |

**Supplementary Table 8.** correlation between microbial function FSCA and ASD symptom.

| index | EC | EC name | ASD symptom measure | Pearson Correlation | p value |
| --- | --- | --- | --- | --- | --- |
| 1 | 2.8.2.22 | aryl-sulfate sulfotransferase | RBSR | -0.500070 | 0.001188 |
| 2 | 3.5.4.13 | dCTP deaminase | RBSR | -0.501642 | 0.001140 |
| 3 | 2.3.1.117 | 2,3,4,5-tetrahydropyridine-2,6-dicarboxylate N-succinyltransferase | CARS | -0.501876 | 0.001133 |
| 4 | 2.3.3.10 | hydroxymethylglutaryl-CoA synthase | ABC | -0.504067 | 0.001069 |
| 5 | 2.7.1.36 | mevalonate kinase | ABC | -0.504282 | 0.001063 |
| 6 | 4.1.1.46 | o-pyrocatechuate decarboxylase | ADOS-SA | -0.505993 | 0.001015 |
| 7 | 2.7.7.59 | [protein-PII] uridylyltransferase | CARS | -0.506344 | 0.001006 |
| 8 | 4.2.1.22 | cystathionine beta-synthase | RBSR | -0.510370 | 0.000902 |
| 9 | 6.3.2.2 | glutamate-cysteine ligase | CARS | -0.514916 | 0.000796 |
| 10 | 1.1.1.88 | hydroxymethylglutaryl-CoA reductase | ABC | -0.517677 | 0.000738 |
| 11 | 6.2.1.22 | [citrate (pro-3S)-lyase] ligase | ABC | -0.522807 | 0.000639 |
| 12 | 2.1.1.172 | 16S rRNA (guanine1207-N2)-methyltransferase | CARS | -0.527377 | 0.000561 |
| 13 | 4.1.1.31 | phosphoenolpyruvate carboxylase | CARS | -0.534938 | 0.000451 |
| 14 | 1.2.7.1 | pyruvate synthase | CARS | 0.531719 | 0.000495 |
| 15 | 2.7.8.35 | UDP-N-acetylglucosamine-decaprenyl-phosphate N-acetylglucosaminephosphotransferase | ADOS-Total | 0.502404 | 0.001117 |
| 16 | 2.7.8.33 | UDP-N-acetylglucosamine-undecaprenyl-phosphate N-acetylglucosaminephosphotransferase | ADOS-Total | 0.502404 | 0.001117 |

## Notes

### Competing Interest Statement

The authors have declared no competing interest.

